# Thermo-Dynamic Flux Balance Analysis, a novel approach to simulate circadian metabolism

**DOI:** 10.64898/2025.12.17.694837

**Authors:** Gian Marco Messa, Peng Liu, Francesco Napolitano, Jesper Tegner, Xin Gao, Valerio Orlando

## Abstract

Circadian metabolism arises from complex, time-dependent interactions across organs, yet experimental characterization in humans remains limited. Current whole-body metabolic models are either too large for dynamic simulation or insufficiently detailed to capture temporal physiology.

We developed a multi-tissue human metabolic reconstruction, *HGEM1*.*19+*, an extensively curated model for dynamic simulation. We introduced Thermo-Dynamic Flux Balance Analysis (tdFBA), a novel formulation that integrates thermodynamic constraints, organ-specific enzyme capacities, solubility limits, osmotic balance and pH buffering. Murine circadian transcriptomics and metabolomics were integrated with human datasets to parameterize temporal dynamics. A genetic algorithm optimized internal parameters under CHOW diet conditions and model performance was evaluated using circadian metabolomics and gene-sage patterns.

The framework recapitulated key metabolic oscillations under CHOW diet and partially reconstructed perturbed states including HFD, BMAL1 knockout, and tissue-specific re-entrainment.

tdFBA enables physiologically grounded, multi-organ dynamic simulations and provides a foundation for exploring human circadian metabolism and perturbation responses.

## 1. Introduction

Circadian metabolism is a central driver of physiological homeostasis (Koronowski and Sassone-Corsi, 2021), yet its dynamics in humans remain largely unexplored due to limited temporal data and the complex interplay between systemic metabolism and environmental cues, highlighting the need for predictive computational models.

In this work, we present the Thermo-Dynamic Flux Balance Analysis (tdFBA), a novel approach to explore the dynamics of complex biological systems *in-silico*. Our method leverages extensive gene expression, proteomics, and metabolomics datasets generated over recent years into a unified framework for system-level modeling.

Due to the nature of our study, we sought to limit the scope of our framework to target time scale around 24/48h to emphasize the circadian phenomena and changes happening within an organism. The circadian rhythm is, in fact, deeply rooted in our physiology and it is fundamental to optimize the metabolic and homeostatic functions of the organism (Asher and Sassone-Corsi, 2015; Reinke and Asher, 2019).

Although the framework can, in principle, simulate indefinitely, its current computational scalability limits its application to longer biological phenomena, such as seasonal rhythms or aging.

To cope with the difficulties of data acquisition and incompleteness for human circadian phenomena (Dallmann et al., 2012), we integrated a vast amount of information comprising gene expression data (Eckel-Mahan et al., 2013; Lonsdale et al., 2013; Masri et al., 2016; Koronowski et al., 2019), metabolomics (Dyar et al., 2018) and proteomics (Bryk and Wiśniewski, 2017), sourcing and adapting both human and murine data.

We decided to focus our efforts in model developing toward the central role of time, embedding it as a core feature of our strategy. Moreover, we sought to strongly integrate in our system several physiological conditions and feedback loops.

Our modeling framework is able to reproduce a dynamic whole-body metabolism (WBM), which can be subject to and tested with any type of perturbation in gene expression and diet.

To achieve this objective, we applied the combination of metabolic networks and Flux Balance Analysis (FBA) (Palsson, 2015). This first principle approach allows us to produce a vast amount of simulated data in a relatively low computational cost, since these simulations can run on commodity hardware.

Most importantly, this approach relies on setting rules and using physical laws that can be explicitly interpreted or modified, derived from publicly available studies and data, making this approach transparent and interpretable, in contrast with pure machine learning methods.

In the literature, several attempts have been proposed to construct a WBM simulation. These efforts generally fall into two contrasting strategies, which we can refer to as “wide” and “tall,” distinguished by the scale of the metabolic model and the algorithms employed.

In the work of Thiele et al. (2020), the authors approach the problem with a “wide” mindset, using a single metabolic network spanning 26 organs and six blood cell types with over 80,000 biochemical reactions. The principal advantage of this model lies in its completeness and resourcefulness. However, this level of complexity introduces substantial limitations from both physiological reconstruction and computational perspectives, particularly with respect to our objectives. Algorithmically, the authors employ Flux Variability Analysis (FVA) (Mahadevan and Schilling, 2003), a variant of FBA that provides a detailed characterization of metabolic capabilities at steady state. While well suited for static analyses, this choice becomes impractical in the dynamic context required for circadian simulations due to the prohibitive computational time and memory demands associated with each time step.

An alternative strategy for studying whole-body metabolism is the “tall” approach, which models a limited number of organs, therefore scaling down the problem to allow even more expensive algorithms. This strategy is exemplified by Martins Conde et al. (2021), who simulate dynamic metabolism using liver, skeletal muscle, and white adipose tissue through dynamic FBA (Mahadevan et al., 2002). In this work, we adopt a similar conceptual approach, but introduce a substantially more detailed representation. Moreover, our algorithmic choice is distinct and novel, designed to balance computational efficiency with the dynamic nature of the problem.

To construct our framework we set several objectives to guide our model and algorithmic choices, with maximizing physiological validity as the top priority.

As the metabolic network backbone, we opted to use HGEM1.19 (Robinson et al., 2020). This model is in continuous development by the authors and the community on GitHub, facilitating access and modification.

Recon3D (Brunk et al., 2018) represents a closely related reconstruction with comparable scope. However, its development and distribution model differ and public updates are not available, to the best of our knowledge. Both models are supersets of Hepatonet1 and similar manually curated models (Gille et al., 2010; Pagliarini et al., 2016).

Nevertheless, since HGEM1.19 is a GEM partially generated automatically, we performed extensive manual curation of the network, generating a new model, which we named *HGEM1*.*19+*. The details of the modifications to the network are in the Methods section with an overview of the final structure of the model shown in Figure 1.

**Figure 1.**
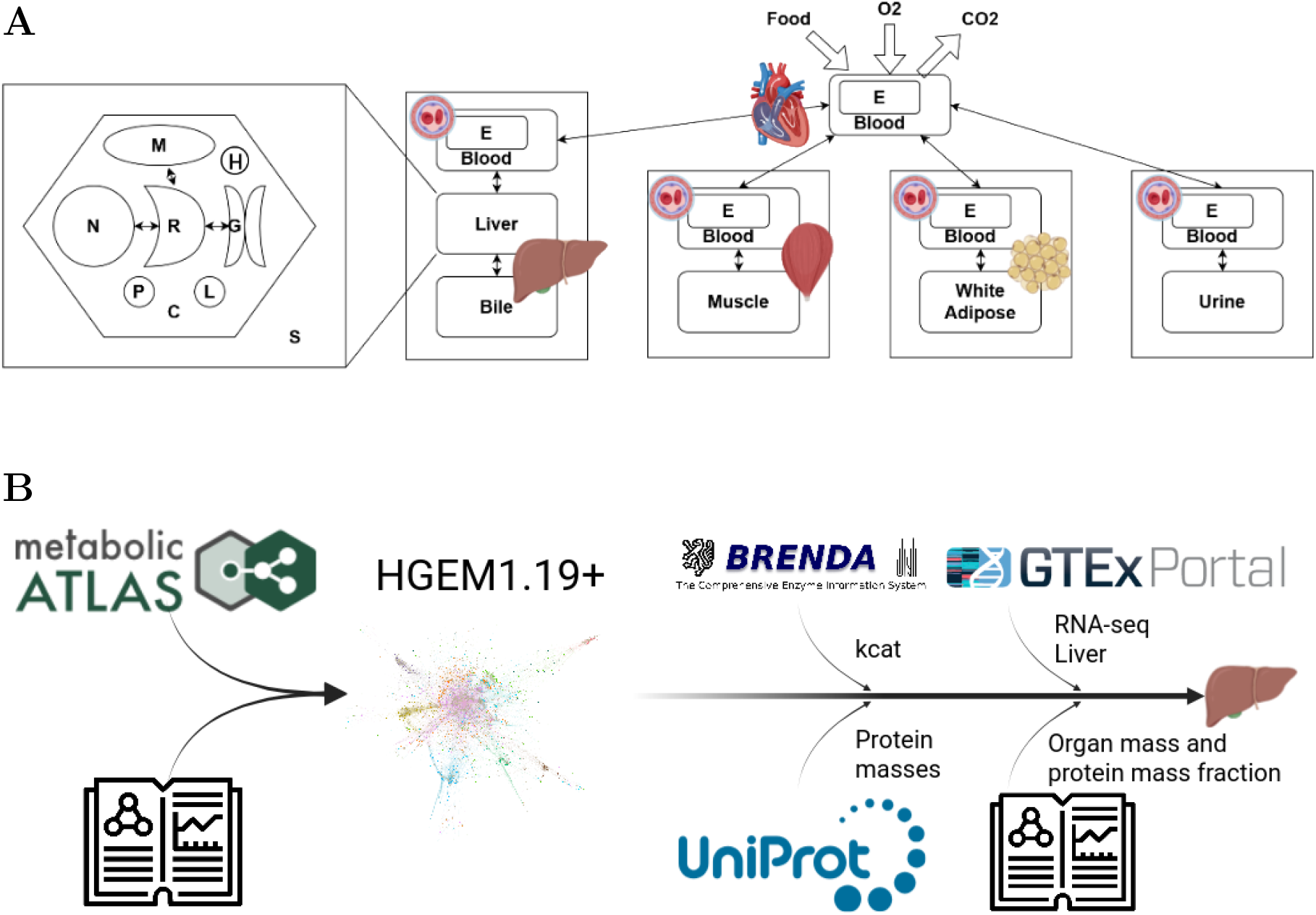
**A**: Schematic of the model structure. Each organ is subdivided in the principal organelles: Cytosol (c), Nucleus (n), Peroxisome (p), Endoplasmic Reticulum (r), Mitochondria (m), Lysosome (l), lipid droplet (h), External Space (s). Each organ, if necessary, is connected to their specific compartments, such as the bile for the liver, and to the blood. The blood, in turn, contains a compartment erythrocyte that is circulated with the blood itself. At each iteration of the algorithm the blood is pooled together and averaged before being replenished and redistributed. **B**: The information flow generating the tissue specific model. The base model HGEM1.19 from the metabolic atlas is manually integrated with literature data generating the HGEM1.19+ model. Then the data from BRENDA, UNIPROT, GTEx and physiological characteristics of the organ extracted from literature are used to constraint the model assign organ specificity.

In this study, we present a novel modeling and algorithmic framework that integrates multi-omics data to generate physiologically realistic, dynamic WBM simulations, in which circadian rhythms emerge naturally from the system. We demonstrate its application to dietary perturbations, such as High Fat versus Chow diets, revealing mechanistic insights into temporal metabolic dynamics at both cellular and systemic levels.

## 2. Results

The main aim of this work consists in developing and validating a WBM simulation framework that will allow for stable and easy to access insights on the WBM in different conditions. To achieve this result we iterated over a very large number of different approaches, algorithm and modeling choices, for a full description please read the Methods section.

To briefly recap our process, we developed Thermo-Dynamic FBA (tdFBA) formulation that intrinsically requires the simulations to be dynamic to avoid NP algorithmic complexity. To make this algorithm work it is required to know a-priori an extremely large number of parameters that define the behavior of the simulations. Some of these parameters can be computed using other software, such as Chemaxon (Chemaxon) and Equilibrator 3.0 (Beber et al., 2022), or extracted form literature, many others are beyond prediction or impossible to determine experimentally. Therefore we set up a genetic algorithm to search for these parameters in such a way that they would be independent from the conditions.

To reach this scope, we start our search on the most optimal conditions for the physiology of the organism, in a CHOW diet system. After achieving reasonable results for the CHOW diet, we move on to test the parameters on the HFD and the other perturbed conditions.

In this preliminary manuscript, we will not go into details about detailed biological interpretations of such results but we will limit ourselves to report the stability and the performance of the simulations with respect to the available data. Also, we will not perform additional validations at this stage, such as enzyme KO to simulate inborn error of metabolism or directly compare our results with already available models.

Moreover, in this instance we will also not make the codebase available on github since the final scope of this project is to develop a platform that can be easily queried by biologist and bioinformaticians alike.

### 2.1. A working CHOW diet multi-tissue model

The first challenge we had to face has been to create a working model that could satisfy the conditions we imposed to the simulations, the ability to perform dynamic simulations and then to be physiologically relevant. As such we decided to utilize a genetic algorithm to optimize the internal parameters of the simulation (Δ*G*^0^, allosteric constants and objective function definition) and so train our model on a real metabolomics dataset (Dyar et al., 2018).

Given the enormous size of the search space for the parameters of the model and the poor convergence guarantees of the genetic algorithm, once we consistently obtained models able to run for over 48*h* in CHOW diet condition, we decided to select the top 30 scoring parameter settings and use the models as an ensemble of predictors to have a more flexible result than a single model.

The proposed simulation is able to partially recapitulate the metabolic profiles of both the CHOW diet, used to estimate the parameters of the model, and the HFD.

From the metabolites identified in the metabolomics experiments of the Atlas of Circadian Metabolism Dyar et al. (2018), we managed to map against the large number of metabolites in the network (over 4000) an average of 57% of them. For the CHOW diet, 41% of the mapped metabolites have positive Pearson correlation with the experimental data (*p*–*value* ≤ 0.05) and 13% of these are circadian. For the HFD, unseen by the model in any circumstances, we have still 41% of positively correlated metabolites, of which 24% are circadian. A more detailed view of this information is in table 1.

**Table 1:** Metabolites mapping. In table are reported the number of metabolites mapped from the model to the experimental data of the Atlas of Ciracadian Metabolism Dyar et al. (2018).

| | | Identified Metabolites | | Positively Correlated ($p\text{-value} \leq 0.05$) | |
| --- | --- | --- | --- | --- | --- |
|  |  | Real \| Matched \| (%) |  | Pearson \| Spearman |  |
| Tissue | Total | Circadian CHOW | Circadian HFD | CHOW (Circadian) | HFD (Circadian) |
| Liver | 306 \| 189 \| 62.1% | 103 \| 51 \| 49.5% | 99 \| 78 \| 78.8% | 87 (20) \| 80 (20) | 85 (32) \| 90 (35) |
| Skeletal Muscle | 543 \| 268 \| 49.4% | 132 \| 38 \| 28.8% | 109 \| 64 \| 58.7% | 96 (9) \| 104 (10) | 109 (25) \| 110 (22) |
| White Adipose Tissue | 184 \| 120 \| 65.2% | 4 \| 1 \| 25% | 11 \| 2 \| 18.2% | 39 (1) \| 37 (1) | 39 (0) \| 32 (0) |
| Serum | 362 \| 183 \| 50.5% | 135 \| 26 \| 19.3% | 75 \| 45 \| 60% | 89 (12) \| 100 (14) | 81 (17) \| 87 (23) |

The data in output from the model and from the experimental data have been post-processed to have similar scales for representation purposes. There are a few key differences between the two datasets, first the experimental data are comprised of only 6 or 7 time-points over 24*h*, while the simulation has at least one time-point every 30*s*. This implies that, in order to be able to compare them, the two time-series have to be projected to the same temporal axis. There are several techniques that can be employed for this task, we chose to interpolate the means of the experimental data and resample them with 1 minute of step, instead for the simulated data we took the average of windows of 1 minute, therefore obtaining the same number of points for the two data. Taking an average over a window for the simulated data also helps with partially smoothing the signal.

Another important point is that as replicates for the simulation, in the constructed ensemble, we took models with only small differences in terms of metabolite concentrations. This causes this replicates to have very low variance compared to the matching experimental data. This and the fact that we can virtually have as many replicates as we can compute, makes the data output of the models incompatible with traditional tools for circadian analysis, such as JTK-cycle Hughes et al. (2010) (even after sub-sampling every 4 hours) reporting an objectively excessive number of significative circadian metabolites with very small p-values. Therefore we decided not to include this analysis in the results and proceed with more traditional approaches. In figure 2 are reported 8 metabolites with very high Pearson and Spearman correlations in both CHOW and HFD for all the four tissues of interest. As shown, the models can follow similar patterns with respect to the experimental data which allows us to be reasonably confident that the numbers in output are not completely unrealistic.

**Figure 2.**
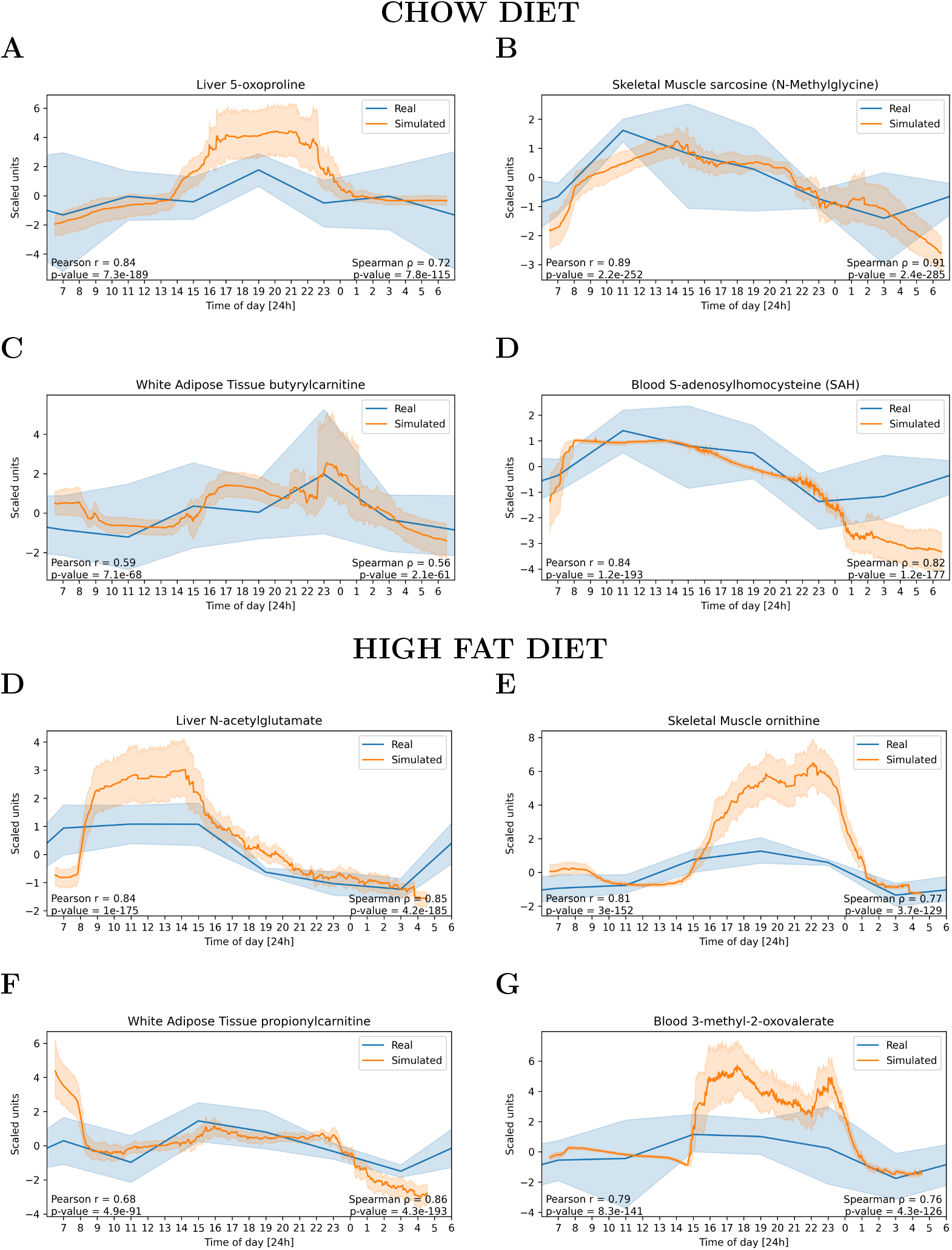
Plot of eight metabolites in CHOW and HFD. The plots show the real (blue) and the simulated (orange) time course data for eight different metabolites in the two conditions of CHOW and HFD. For these metabolites the correlation coefficients are extremely high and with a very low p-value, in fact the simulation is able to reconstruct profiles which are able to reconstruct the behavior of the real data with very good approximation. It is possible to notice the different sampling frequency of the two datasets by the level of high frequency noise present in the simulated data compared to the real ones.

Moreover, we decided to also interrogate the information carried by the fluxes inside the model. Since we have a direct mapping between reactions and genes, taking the absolute value of the fluxes passing through each reaction and assigning it to the associated genes, we can obtain a proxy for the gene usage at different time-steps inside the simulation. To do so we had to reduce and subsample the outputs to the 6 principal time-points, at ZT from 0 to 20, to make the computations of the statistics last a reasonable time.

To have a better understanding of the behavior of the model in different conditions, we computed the correlation in time of the gene usage values for all tissues in the CHOW and HFD simulations, shown in figure 3. The CHOW diet simulation figure 3A behaves as expected and in the heatmap it is possible to notice a few trends. First of all the squares around the diagonal values all show very high correlation due to the fact that these are all replicates of the same time-point, this means that the model, even with different parameters, is able to behave consistently and does not diverge too much. The pattern also suggests that the model is stable in its performances, at least up to the values shown. It is also possible to notice that there is a strong divergence between the initial state and the following time-points, this is due to the fact that simulation has a set starting configuration which is the same for all possible combinations of conditions, therefore the divergence from the first step is expected. Nevertheless, the correlation between subsequent time-points seems to remain fairly stable, reflecting the convergence of the simulation toward a stable state.

**Figure 3.**
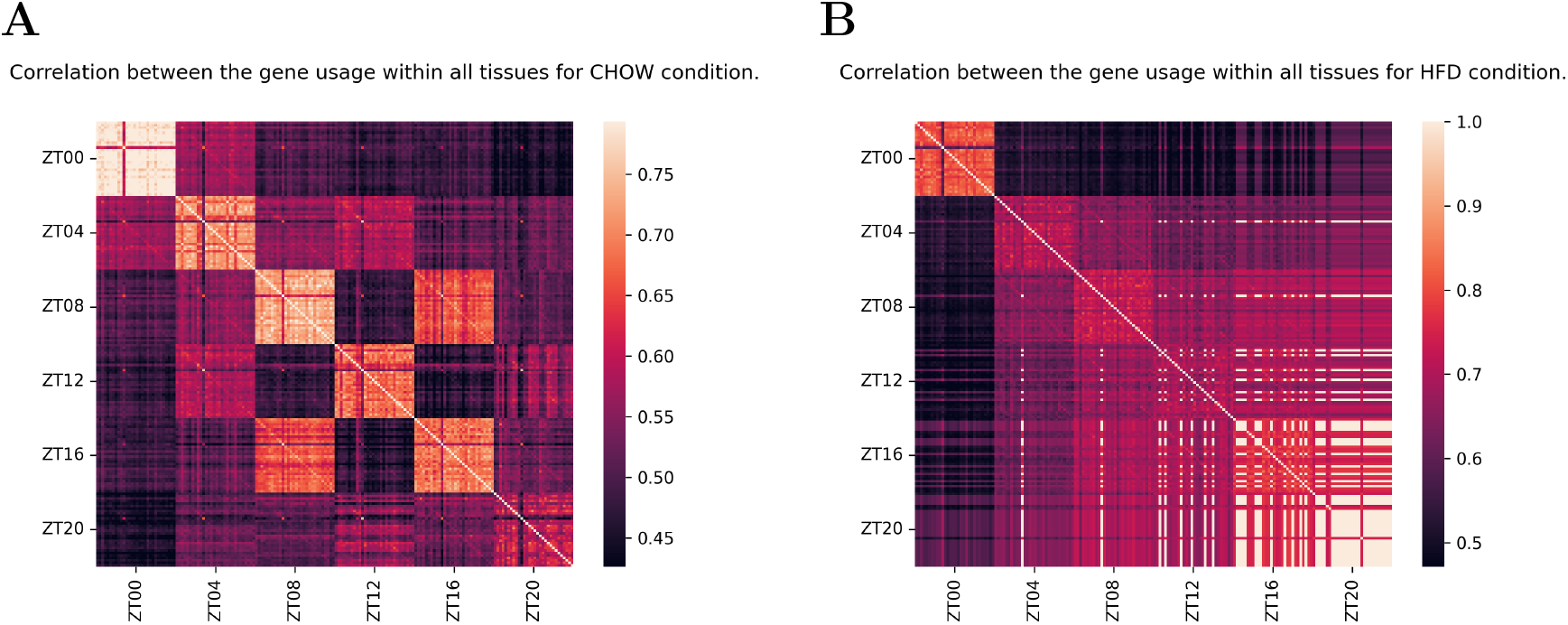
CHOW and HFD heatmaps. The two heatmaps show the correlation between the simulations over the six main simulated time-steps. CHOW diet simulations arrive all at the desired endpoint of ZT20 instead some of the HFD fail before being able to reach a conclusion. The trends of correlations in the heatmaps show consistency between the members of the ensemble and a divergence from the starting conditions with along the time axis, still keeping very high correlation values, as expected.

Regarding the HFD simulation figure 3B, it lacks the strong correlation observed in the CHOW but it also shows the diverging trend from the initial conditions. To be noted is that the correlation around the diagonal is still very high. In this simulation it is possible to observe the failure of more and more replicates which, due to the new gene expression and dietary conditions, are unable to remain stable with the current parameter settings. Fortunately not all models fail in this context and this implies that further training and a new training strategy are required to find the parameters to satisfy a larger variety of conditions. This means that, although promising, the simulation is still not polished enough to reproduce the long term conditions required to ensure a reliable reconstruction in the view of a circadian and periodic output.

### 2.2. Enrichment analysis on model fluxes

Even though these models are not perfect, they can still partially reflect the overall behavior of the perturbed system. To explore this point of view we used the data produced with input the wild type, bmal-KO, liver-RE, muscle-RE and double-RE gene expression data from Koronowski et al. (2019); Smith et al. (2023). The diet given to the model is always the CHOW diet from the previous experiment for all conditions. In figure 4 it is shown the same kind of heatmaps as in figure 3. Unfortunately in this case, for all conditions and for all replicates, the simulation is unable to reach the ZT20, this may be due to a slightly different input data in terms of gene expression, nevertheless, up to the point where the simulations fail, the model behaves consistently with expectations. With high correlation in the area around the diagonal and a progressive reduction of correlation between the first and the following time-steps.

**Figure 4.**
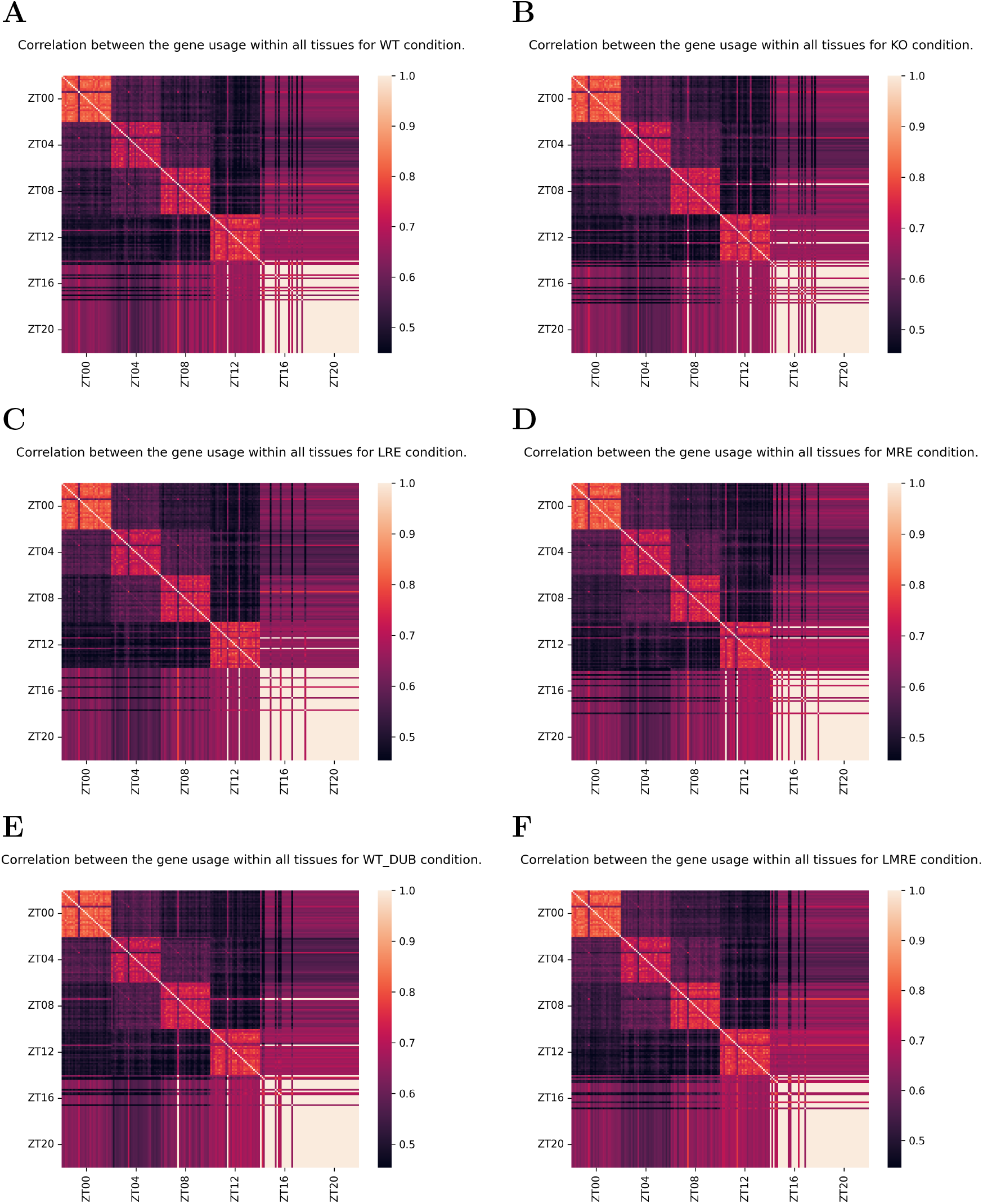
Bmal-KO heatmaps. A) Wild type, B) bmal-KO, C) liver-RE, D) muscle-RE, B) bmal-KO (double RE condition) and F) double-RE. The heatmaps, which show the failure of most of the simulations at ZT16 and all of them at ZT20, demonstrate consistency between themselves and the CHOW and HFD simulations, having very similar trends at level of ensembles and inside the replicate groups.

For these data we decided to apply enrichment analysis on the gene usage in the network. Leveraging the GSEA algorithm Subramanian et al. (2005) and the gene usage extracted from the simulation up to ZT16, we produced two different kind of outputs that may give some information about the behavior of the system and if it is sensible enough to actually reconstruct the physiological metabolism. These particular enrichments are performed without taking time into account.

In figure 5 it is shown the comparative enrichment between 6 different conditions, specifically in muscle and liver between their wild types, their RE and the double RE condition.

**Figure 5.**
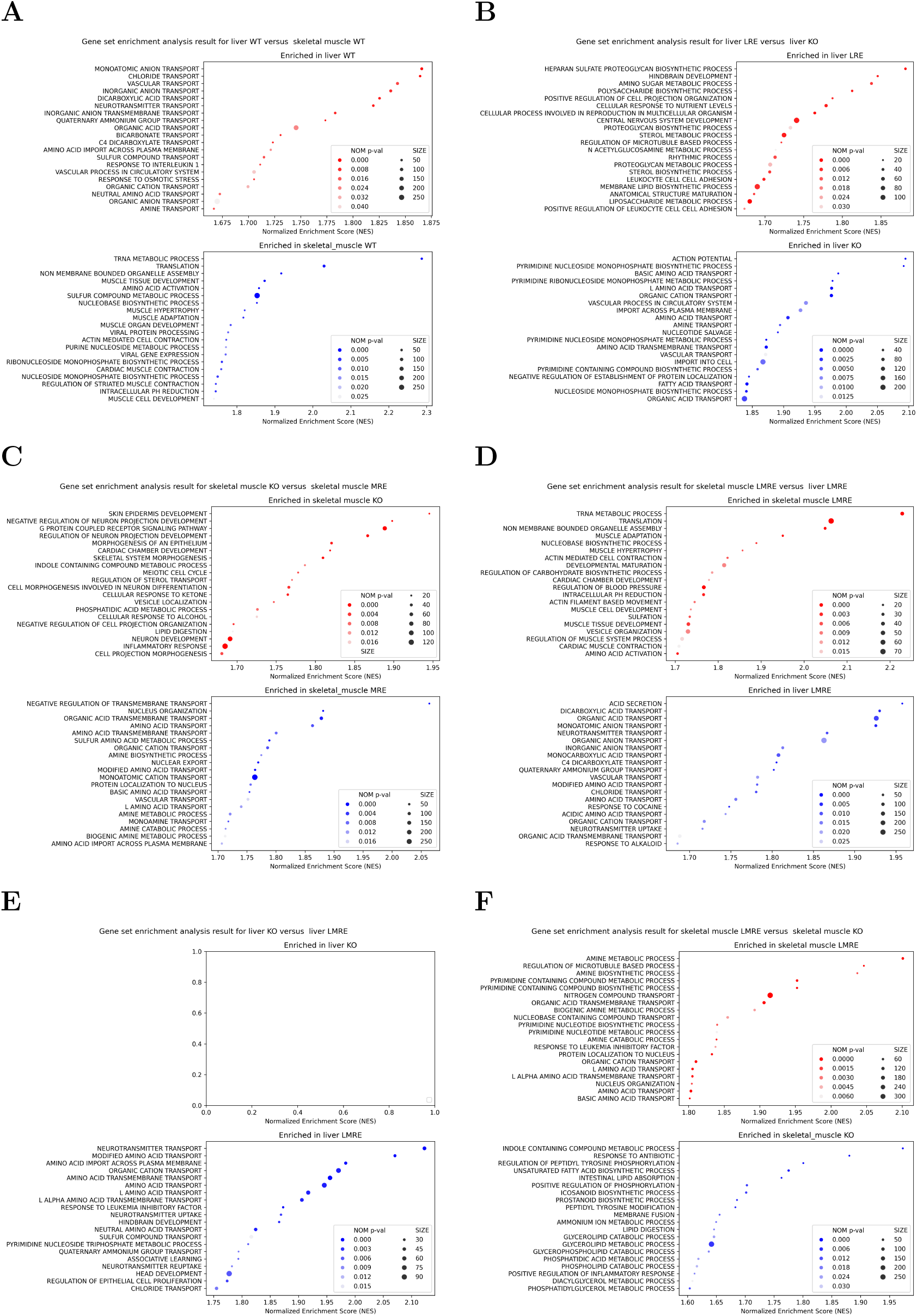
GSEA on liver and muscle for the different conditions of wild type, Bmal-KO, liver-RE, muscle-RE and double-RE. The output of the simulations are compared in the 6 panels of the figure showing the differences between the two organs (A,D) or the same organ (B,C,E,F).

The output of the GSEA shows a very wide variety of terms, sometimes unrelated to metabolism, but the majority, as expected, is driven toward this subset of the ontology. In panel A there is the comparison between the wild types of liver and muscle where the liver shows a prominent activity of metabolic transport with respect to the other tissue which, instead, shows enrichment in specific pathways related to its functions.

In panel B it is possible to notice that the transport activity of the liver is less prominent in its RE condition then in its KO, and the RE is mainly enriched in large molecules and fatty related processes. It is interesting that the liver RE condition shows also enrichment in the term *RHYTHMIC PROCESS* which is a promising result.

The other conditions follow roughly the same patterns, unfortunately, without anything of particular interest that catches the eye except for the comparison between the double RE and the KO condition in the liver showing no significative enriched terms in the KO compared to the RE.

Finally we compared the number of common enriched terms of the different conditions between the various ZT times, shown in figure 6. The Venn diagrams show the overlap between the terms enriched following different peak profiles at 5 different time steps (due to the previous correlation analysis we removed the ZT20). The expected outcome is that the RE conditions are able to recover the KO conditions overlapping a larger number of terms with the WT. Most often in these simulation it is not the case, nevertheless the uniformity of most of the diagrams hints toward the possibility that a stronger bias (that is most likely coming from the unreliability of the parameters) is covering the information regarding the reestablishment of a more similar phenotype between WT and RE conditions.

**Figure 6.**
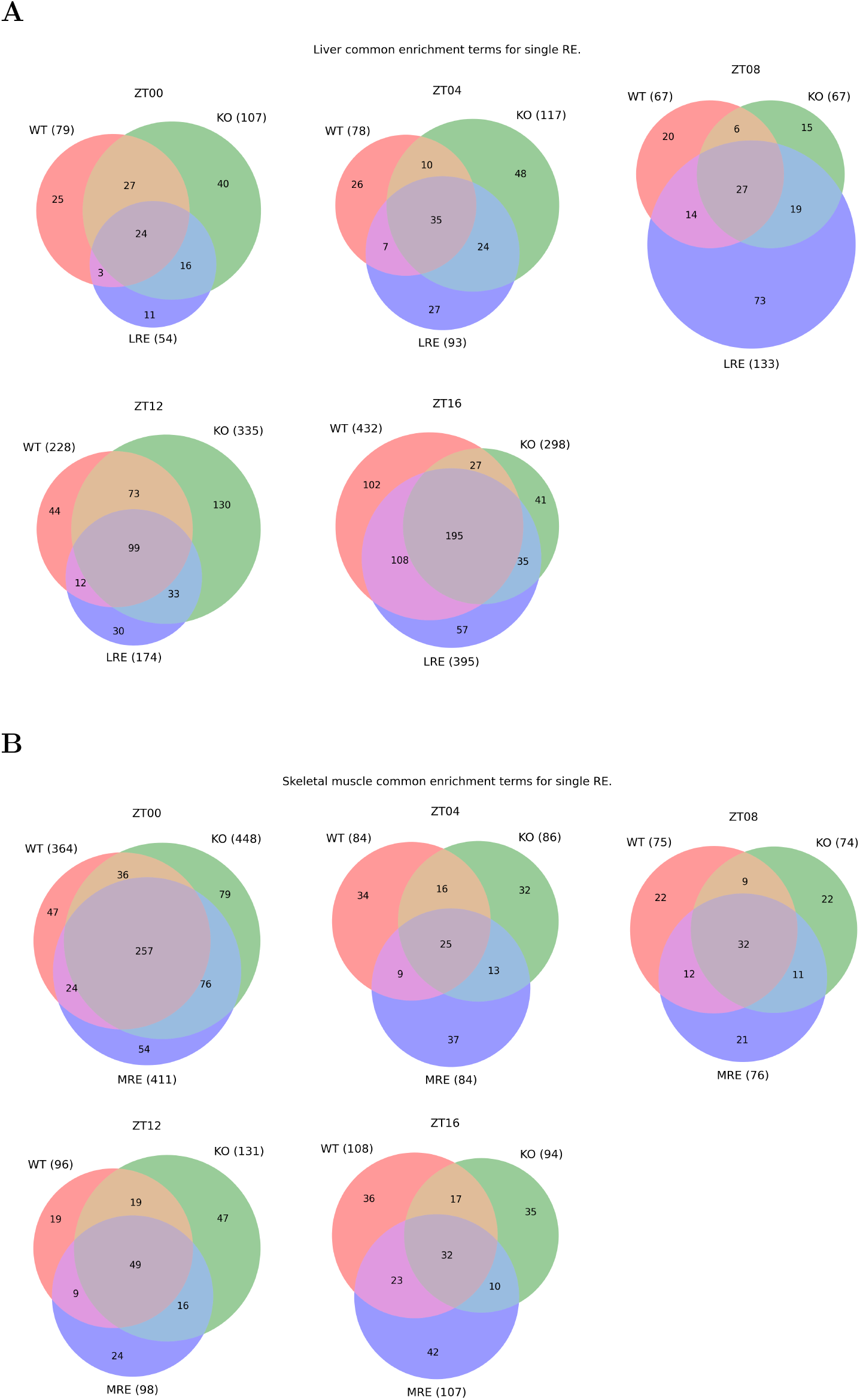
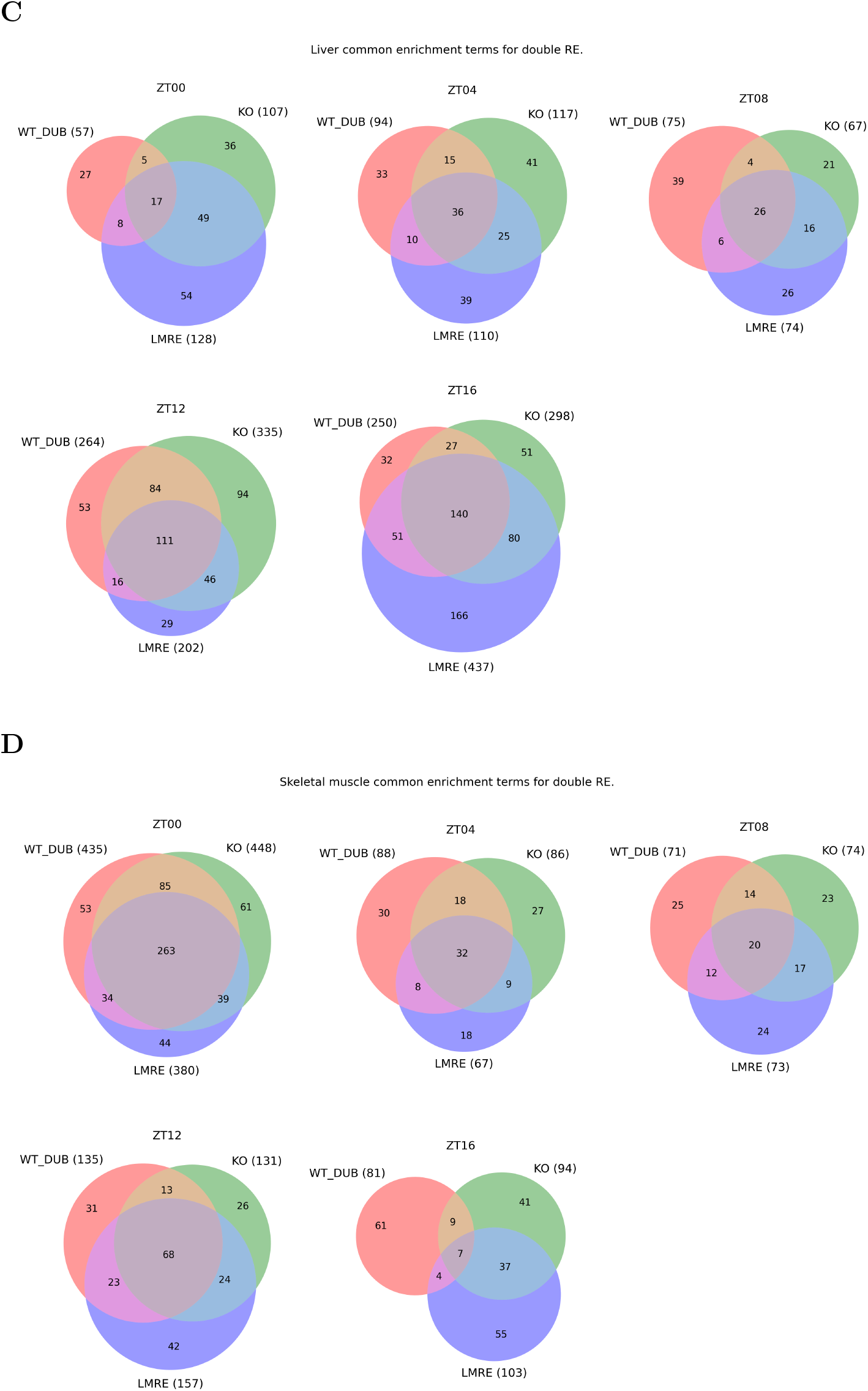
Venn diagram on liver and muscle for the different conditions of A) liver-RE, B) muscle-RE, C) liver-doubleRE, D) muscle-doubleRE. This enrichment is made for the peak phase of the gene usage (either positive or negative) using a continuous phenotype profile in the GSEA software, the diagrams show the number of common significant terms for each sub condition of the experiment with the relevant wild type.

## 3. Conclusions

We built a model and a framework gathering a very large knowledge-base, integrating several scientific disciplines under the same hood in the attempt to create a physiological representation of a human being from the metabolic standpoint.

The model integrates data from human and mouse to overcome the limitations of requiring large amount of data on different tissues of human for several time-points. The model is built with an eye toward dynamic simulations, with the purpose of obtaining emergent oscillatory circadian behaviors without imposing them forcibly or including differential equations.

The model is able to reconstruct acceptably a dietary condition of CHOW diet with wild type gene expression, it also is able to partially reconstruct the behavior of the biological system in perturbed states, with better performances upon dietary stress than the KO of the circadian molecular clock. Unfortunately there are no software made to analyze these kind of synthetic biological data which are different from the sampled data in distribution and unit of measure, therefore a set of proper analysis tools will be developed to extract relevant biological information from these models and make them able to make inference on otherwise unexplorable biological conditions.

For the future it also in development a web interface to submit gene expression profiles and diets to obtain a dynamic metabolic simulation with information about metabolite, genes and reaction fluxes.

## 4. Materials and methods

Moving on from a stationary approach using the tFBA and Hepatonet, to achieve the target of building a model for circadian metabolism, the first thing to do is to characterize the simulation process in a physiological way. One of the major drawbacks of metabolic networks and FBA is the lack of specific information regarding the system, indeed the FBA variants already published tend to underestimate the importance of phenomena such as osmotic balance and how the different organs are connected.

### 4.1. Organ characterization

Our modeling approach has two key requirements: first it has to be dynamic to be able to recapitulate circadian metabolism; second it has to be multi-tissue, representing the interactions between the organs of the system, since it is fundamental in synchronizing the tasks that each organ performs. A multicellular organism does not work if there are some of the major ones missing.

To do so we have to define how the metabolic sub-network is extrapolated from the GEM, which is a generic representation. For several years there have been published tools to create a specialized network from a GEM, such as FASTCORE (Pacheco and Sauter, 2018). Unfortunately these methods irreversibly prune the network to create the specialized networks and we require the entire network to be available to be able to change the parameters of the algorithm based on the changing gene expression. The approach described here acts at the optimization problem level instead of the network itself and leverages the fact that presolve methods of linear program optimizers are extremely efficient to avoid preventively pruning the network.

#### 4.1.1. Tissue specific reconstruction

Our method to specialize the generic GEM is based on the characteristics of the tissue itself, either coming from gene expression data or from other sources. To softly constraint the network to reflect the gene expression we estimate and act on the *V*_*max*_ and *V*_*min*_ (both called *V*_*max*_ from now on for simplicity) of the reaction boundaries. The estimation of the *V*_*max*_ relies on gene expression data or protein abundance from proteomics as a proxy for the real quantities of the enzymes in the tissues. The usage of gene expression data, for the most part, is a critical approximation that has to be done since the availability of proteomics data is limited in terms of data abundance and number of proteins identified.

Combined with the estimate of protein abundance, the *V*_*max*_ definition relies also on the turnover numbers of the enzymes involved, which is the maximum amount of product they can handle in optimal conditions per second. Therefore it is imperative to properly map the information available in the metabolic network to the most common databases to fetch the data required which is done leveraging the gene-reaction rules (GRR) assigned to each reaction.

#### 4.1.2. Protein abundance estimation

After mapping the gene expression and proteomics data to the model, to derive the *V*_*max*_ it is necessary to estimate the protein abundance and the turnover number of the enzyme for that particular substrate.

Starting with the abundance, we adopted a simple practical model with the assumptions that 1) protein turnover (rate at which a protein is degraded) is constant and the same for all proteins, 2) protein synthesis rate is constant and the same for all proteins, 3) proteomics and gene expression data are linearly correlated with the abundance of proteins in the system. Aware that all three assumptions are wrong, they become a necessary evil in order to be able to integrate as much information as possible into the model without developing a protein synthesis and degradation model itself.

Then we proceeded with the following steps: we mapped all the available gene expression data and proteomics data to UniProt and from there we extracted the average mass of the synthesized protein; from the gene expression data, with values in transcripts per million (TPM), we selected the coding genes and normalized the values to have their sum equal to 1; each re-scaled value is then divided by their respective protein mass; finally these values are multiplied by the estimated total protein mass of the tissue taken from literature (Stroh et al., 2021). This gives a value in *mol*/*kg* ready to be properly integrated in the model.

Regarding proteomics data (Bryk and Wiśniewski, 2017), we required them to estimate the abundance of proteins in red blood cells, they were given already in concentration, so it was necessary only to re-scale them in the proper units.

#### 4.1.3. Enzyme turnover number

Given that FBA is not an algorithm that allows modeling unsteady but only steady states, we did not incorporate kinetics in the model, but we still require the *V*_*max*_ of reaction. This value is the product between the abundance of the enzyme and the enzyme turnover number *k*_*cat*_. To estimate this constants we mapped the reactions’ EC numbers to the BRENDA database (Chang et al., 2021) which contains a very large amount of information associated with each reaction.

The mapping of EC numbers is non-trivial, it takes into account deprecated numbers present in the metabolic network and some of them had to be double checked manually. Moreover, in BRENDA are reported the experiments used to calculate the *k*_*cat*_ coefficients, the information includes the species, the vector used to express the enzyme and if the enzyme is mutated or wild type. Only the values associated with the fields “homo-sapiens”, non “mutant”, non “truncated” and with exact substrate were considered acceptable, unknown values are marked with a “-999”.

This process outputs a list of values for each substrate of the enzyme and after observing the results it became clear that some entries were filed inconsistently with respect to the units of measurement. Indeed, sometimes, a value is reported in [1/*h*] instead of [1/*s*], therefore, at processing time, a heuristic checks if the value is an outlier to the distribution (testing if it is greater then 1000 times the smallest in the list) and divides it by 3600, accordingly.

Then for each reaction with associated EC number, it is taken the value of the *k*_*cat*_ extracted and associated with the substrates involved and the final value of *k*_*cat*_ for the reaction is the average.

To finally compute the *V*_*max*_, the values of abundance and the values of *k*_*cat*_ are multiplied with each other and scaled to the proper units of measure.

#### 4.1.4. Organelle characterization

The model we built exploits the sub-cellular compartmentalization already present in HGEM1 with the most common sub-cellular compartments available. We started modeling using HGEM1.10 while, in the update model HGEM1.19 the letters associated with the compartments were changed to be in line with RECON3D. It was our decision, instead, to keep the previous labeling, finding it more readable.

Getting the correct units of measure for this kind of modeling becomes paramount to achieve a physiological simulation of the system. Since we are describing a dynamic system, we have to specify that: the fluxes are measured in [*mmol* · *h*^−1^], the concentrations are in [*mmol* · *kg*^−1^], masses are in [*kg*] and time is in [*h*]. An exception is done for the set of exchange reactions for bile, blood and red blood cells that are expressed in 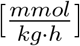 and this depends on the mathematical way in which these compartments are treated.

This choice of units implies the need to estimate the mass proportion of the compartments. This is required to be able to update the concentration of metabolites in time in a correct way across the membranes. Therefore a research in literature (Jiang et al., 2021; Young and Egginton, 2009) and some reasonable assumption brought the proportion described in table 2. These measures of allometry are conducted with cryo-EM technology and, recently, more data is being produced, such as (Parlakgül et al., 2022) (which contains both normal and obese hepatocytes), which was not available at the time of model construction. Nevertheless, with more precise data, it would be possible to represent the tissues with even more details and fidelity.

**Table 2:**
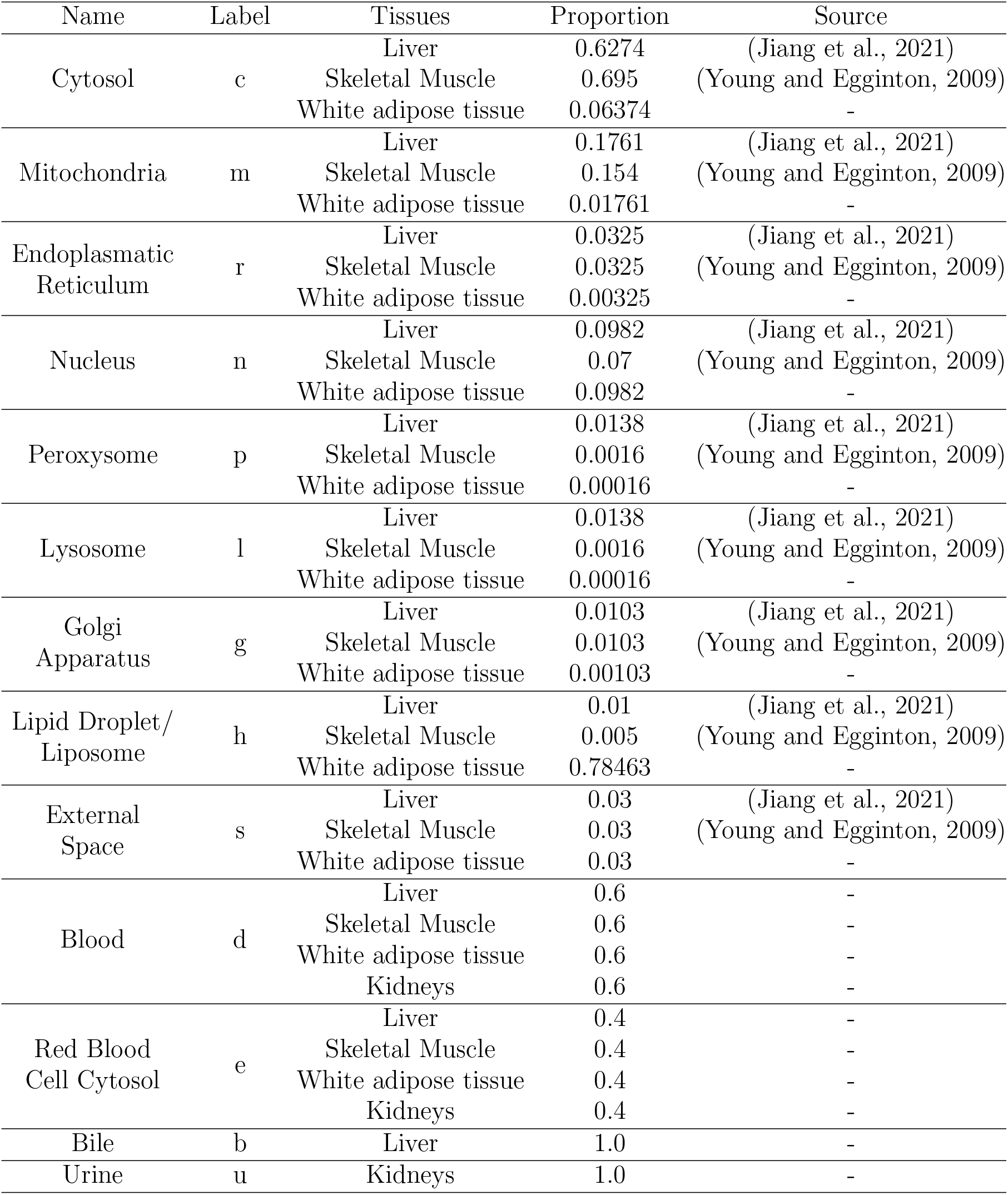
Organelle list and availability to the different tissues. The values are extracted from sources (Jiang et al., 2021; Young and Egginton, 2009) when available. For the liver and the skeletal muscle the data are incomplete and the information has been filled with reasonable values. Regarding the white adipose tissue it was not found any source for this kind of information, therefore the values are simply the once for the liver divided by ten, except for the liposome which is the complimentary of the sum of the other organelle sizes. Blood and RBC are an healthy hematocrit, bile and urine are just placeholders.

To be noted, mitochondria are modeled as a single compartment, which is the mitochondrial matrix. This is because the inter-membrane space of the organelle is extremely porous and allows small molecules and small proteins (*MW <* 10*kDa*) to pass through (Rimessi et al., 2008), so, in the model, it is considered as cytosol.

The endoplasmatic reticulum represents the rough one, the smooth one and the nuclear double membrane, since it is de-facto a single compartment.

The lipid droplet/liposome is the fat storage compartment, it is present in different proportions in all three major tissues and it was non present in the original HGEM1.

The external space is the space between cells and it is put between the tissue and the blood system.

Blood, red blood cells, bile and urine are special compartments, treated differently as explained in the upcoming sections.

Regarding all the the organelles, the model is unable to represent the continuous dynamic changes of proportion, shape, contact, number and so on, so they are treated as static boxes.

### 4.2. Liver, skeletal muscle, white adipose tissue, kidneys

In this section we will describe the organs of interest in the details necessary for the modeling process and their interconnections in the body via the blood stream. Although of fundamental importance for the organism, we decided to omit the lymphatic system which would be extremely complex to implement and would be incompatible with some modeling choices that we had to make.

All the data presented are to be referred to an average human male of 70kg and we assume a total blood flow of 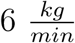.

#### 4.2.1. Liver

The liver is the most chemically active organ in the body, it weights around 1.5 *kg*. It is irrorated by the portal vein and the hepatic artery which, in total, carry around 1.5 − 1.9 *kg*/*min* of blood, being it around the 30% of the entire blood flow available in the circulatory system. The liver has a cell density of 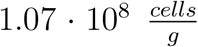, this value is required to estimate the number of nuclei in the tissue.

The blood arriving from the portal vein is full with nutrients coming from the intestines but it is poor in oxygen (75% saturation). The hepatic artery compensates for it coming directly from the principal circulation, nevertheless the liver is always in a slightly hypoxic state, especially the cells further away from the capillaries. We decided not to include the portal system in the model since the digestive system is not included. The existence of the portal circulation also implies the concept of first metabolism, meaning that water soluble molecules are shipped through the portal vein directly to the liver for further processing, instead insoluble molecules, such as fats, are delivered in the subclavian veins by the lymphatic system and are distributed in the rest of the body by the blood stream.

The liver produces on average 620 *mL*/*d* of bile (Esteller, 2008) excreted in the biliary ducts and then in the gallbladder. It consumes around 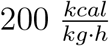 (Wang et al., 2010) 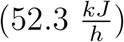, table 3. It is the only producer and the major recycler of albumin, the most abundant protein the blood. It is responsible for the degradation of the HEME groups from hemoglobin. It is able to store about 100 *g* of glycogen, with the second largest capacity in the body (Wasserman, 2009). The liver is composed of around 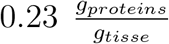. To be noted is that it is very hard to query and get meaningful results for this kind of information since the majority of the results both in literature and just on the web, for example for the protein fraction, are flooded with dietary information and nutritional values of food.

**Table 3:** Specific power consumption by organ/tissue.

| Organ/tissue | Elia's Ki value (Elia, 1992) | 21–30y (young) | 31–50y (middle-age) | 51–73y (old) |
| --- | --- | --- | --- | --- |
| | $\text{kcal} \cdot \text{kg}^{-1} \cdot \text{day}^{-1}$ | $\text{kcal} \cdot \text{kg}^{-1} \cdot \text{day}^{-1}$ | $\text{kcal} \cdot \text{kg}^{-1} \cdot \text{day}^{-1}$ | $\text{kcal} \cdot \text{kg}^{-1} \cdot \text{day}^{-1}$ |
| Liver | 200 | 202 (199, 205) | 199 (197, 202) | 194 (191, 197) |
| Kidneys | 440 | 443 (437, 450) | 438 (432, 444) | 426 (420, 433) |
| Skeletal Muscle | 13 | 13.1 (12.9, 13.3) | 12.9 (12.8, 13.1) | 12.6 (12.4, 12.8) |
| Adipose Tissue | 4.5 | 4.54 (4.47, 4.60) | 4.48 (4.42, 4.54) | 4.36 (4.29, 4.43) |
| Residual | 12 | 12.1 (11.9, 12.3) | 12.0 (11.8, 12.1) | 11.6 (11.4, 11.8) |

#### 4.2.2. Skeletal Muscle

The skeletal muscles account for a very large fraction of the body weight, around 40 % for a total mass or 28 *kg* in our average male. Red muscle cells are syncytia with several thousands nuclei (3,000) per fiber (Snijders et al., 2020). To model human muscles we referred to allometric data from the biceps, we considered 253,000 fibers in the biceps, with a mass density of 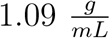, a volume of 160 *mL* and weight of 0.1744 *kg*(Klein et al., 2003; Adkins et al., 2021). The muscle nuclear density is then normalized to the value of 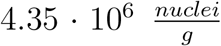 and the organelles usually lie on the side of the fibers. These values from the biceps are considered the same for the entire musculature of the body.

The muscles are irrorated by 3 *to* 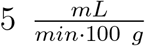 of blood in resting state and can reach up to 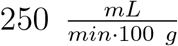 (LEMONS and DOWNEY, 1994). We considered a resting value of 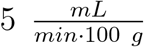 for a total blood flow of 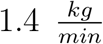. The muscles at rest consumes 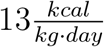 (Wang et al., 2010) (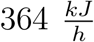, table 3) and to this value we add 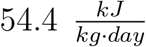 (Cameron et al., 2002) to account for the contraction activity, also we require this additional amount to be consumed exclusively from ATP hydrolysis. The muscles are the main containers of glycogen, accounting for 500 *g* of it in the body (Wasserman, 2009). They have a very peculiar structure concerning the endoplasmatic reticulum, the name is conventionally changed in sarcoplasmatic reticulum and it has a different structure then other cell types, enveloping the muscular fibers like a reticulated glove. The sarcoplasmatic reticulum is a reservoir of calcium ions (*Ca*^2+^), expressing high values of calsequestrin (Sanchez et al., 2012) and the SERCA pump (Murphy et al., 2009). The calcium is released in the cytosol upon stimulation from the action potential triggered by the neural impulses. This process is unfortunately too fast to model with our tools so the SERCA and the calsequestrin are used only to keep the homeostasis of calcium in check against a small leaking flux.

#### 4.2.3. White adipose tissue

The white adipose tissue is the larges, long term, energy storage tissue in the body where fat and non-polar molecules are accumulated. On average fat tissue accounts for 19 *kg* of weight (Dadson et al., 2017), it has a cell density of 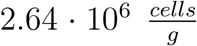. the fat is irrorated by 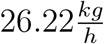 of blood (Jansson et al., 1996), the power consumption is 14.9 *kJ*/*h* (Wang et al., 2010), table 3. The white adipose tissue is not particularly metabolically active. We consider, for simplicity, that there is only one type of adipose tissue and we assume it is represented by the subcutaneus.

#### 4.2.4. Kidneys

Our model does not contain a sub-model for the kidneys, instead this is a placeholder organ consisting of only blood, red blood cells and urine, to allow the disposal of blood molecules in excess, a fundamental process required by the setup to work properly. The data of excretion are extracted from HMDB (Wishart et al., 2018) like in (Martins Conde et al., 2021). The values are normalized according to the units of measure of our model. The kidneys filter on average 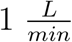 of blood (Sehdev., 2023), without considering the blood required to irrorate the tissue.

### 4.3. Bile and urine, blood and red blood cells

These four compartments require special attention since they are the only ones that are open to the external world or behave as connector between the tissues.

Bile and urine have excretory work. The liver is the only organ which communicates with the bile from the cytoplasm and secretes the relevant molecules keeping physiological concentrations (Hall and Guyton, 2006; Boyer, 2013). Since our model is essentially 0 dimensional in space, we did not consider the polarization of the hepatocytes, therefore there is no distinction between the apical part of the cells and the rest of the membrane. Moreover we did not consider the different cell types that interact with this compartment, such as cholangiocytes.

The urine compartment is directly accessed from the blood and only in the kidneys, the molecules transported in the urine compartment are simply removed from the system in the amounts found in HMDB (Wishart et al., 2018) if present in sufficient amount in the blood. For the bile we keep track of the concentration of the metabolites in the compartment, since it is the major driver of excretion and it is simple enough to model, instead for the urine we do not because the nephron is a too complex structure to be modeled with this means.

Blood and red blood cells are also modeled in a different way then the other compartments, first of all they do not have associated a volume or a mass but only a flow, the blood compartment communicates with the external spaces of the tissues and the red blood cell compartment communicates only with the blood. This allows for the relevant reactions to happen in the red blood cells and the blood (which is then more properly the serum) is a communication and transport compartment. During the simulation the nutrients are directly inserted in the blood stream and the hemoglobin in the red blood cells saturated. The *CO*_2_ removed, without an explicit simulation of lungs or intestines.

### 4.4. Circadian metabolism

As stated previously, we extracted and mapped gene expression data to characterize the organs of the model. To achieve so we acted on two subsequent layers. First, from the GTEx project database (Lonsdale et al., 2013), we selected the best male candidate to be representative of the human species, the criteria for our choice were based on the availability of all the tissues of interest from the same donor and the high quality of the RNA extracted. We do not have access to detailed metadata for this dataset since this information requires the submission of a project to the GTEx consortium. In the future we will definitely gather this information and leverage it to further perfect the simulation. The candidate subject is identified as “OIZF” and the liver, skeletal muscle and white adipose tissue have, respectively, the following identifiers: GTEX-OIZF-0826-SM-3MJGO; GTEX-OIZF-1626-SM-2YUNE; GTEX-OIZF-0226-SM-E9TI3.

This sample represents a single, unknown, time-point for the reconstruction of a circadian profile. To best exploit the data available to us we decided then to integrate the large amount of murine data available from Eckel-Mahan et al. (2013), (Smith et al., 2022) and (Xin et al., 2022), plus some data produced in house.

### 4.5. Murine data integration

#### 4.5.1. Gene expression

To connect gene expression data between mouse and human we generated an orthology table querying ensemble.

With this mapping done, we then extracted from the datasets availabe from Eckel-Mahan et al. (2013) the gene expression for the liver in normal chow diet (CHOW) and high fat diet (HFD), which have been produced with microarray technology. This implies that the dynamic range of the data is much tighter than RNA-seq next generation sequencing by Illumina, spanning only 3-4 orders of magnitude instead of 5-6 (Zhao et al., 2014). So, to compensate for these different distributions, we decided to take two to the power of the reported data and perform geometric mean normalization and standardization with a Z-score. Then we applied another exponentiation calculating *e* (Euler’s number) to the power of the result. Therefore we obtained the profile for the liver in the two different dietary conditions.

From in-house, still unpublished, experiments we processed the data for the skeletal muscles, being sequenced with RNA-seq technology, we did not perform the first exponentiation and started from the data in TPM. Since there are many zeros in this dataset, we added a very small noise sampled from a Gaussian distribution with mean 0.05 and standard deviation 0.05, much smaller then the relevant values in the data, in order to circumvent division by zero errors. We performed geometric mean normalization and standardization with a Z-score and applied exponentiation again getting *e* to the power of the result.

We repeated the same procedure used for the skeletal muscle for the white adipose tissue RNA-seq data from Xin et al. (2022) renormalizing them to TPM before starting.

The time-point in the circadian field are identified as ZT, standing for Zeitgeber, where ZT0 is the beginning of the light and ZT12 the beginning of darkness. We have then 7 time-points for the liver gene expression data from ZT0 to ZT24 sampled at intervals of 4 hours of which we used the ZT24 as replicates for ZT0, we have 6 time-points for the skeletal muscle from ZT0 to ZT20 sampled at intervals of 4 hours and we have 2 time-points for the white adipose tissue sampled at ZT0 and ZT12.

We also have access to the datasets for the bmal-KO experiments of (Koronowski et al., 2019) and (Smith et al., 2023) which were processed in the same way as the skeletal muscle, being produced with RNA-seq technology.

For the actual integration of the two mammalian datasets, the data from GTEx is processed as described in section 4.1.1 to transform it in the *V*_*max*_ base values, then the murine data with two or more time-points is used as a modulator for the expression in human in time, following the map of the orthologue genes, extracted as described previously. Since human and mouse have different active time with respect to the light/darkness cycle (mice are nocturnal animals), we shifted the profile by delaying the clock of 5 hours (*ZT*0 → 7 *pm* and so on), in this way we can synchronize the beginning of daylight for mouse with sundown for human, overlapping the active and resting phases. We then computed the cubic spline passing through the average of each time-point enforcing periodicity. In this way we have a mathematical structure that we can query as needed for each possible time-point. In figure 7 there is a representation at one time-point (ZT7) of the resulting *V*_*max*_ values, calculated with this process, for a visual clue of the relative values between the tissues.

**Figure 7.**
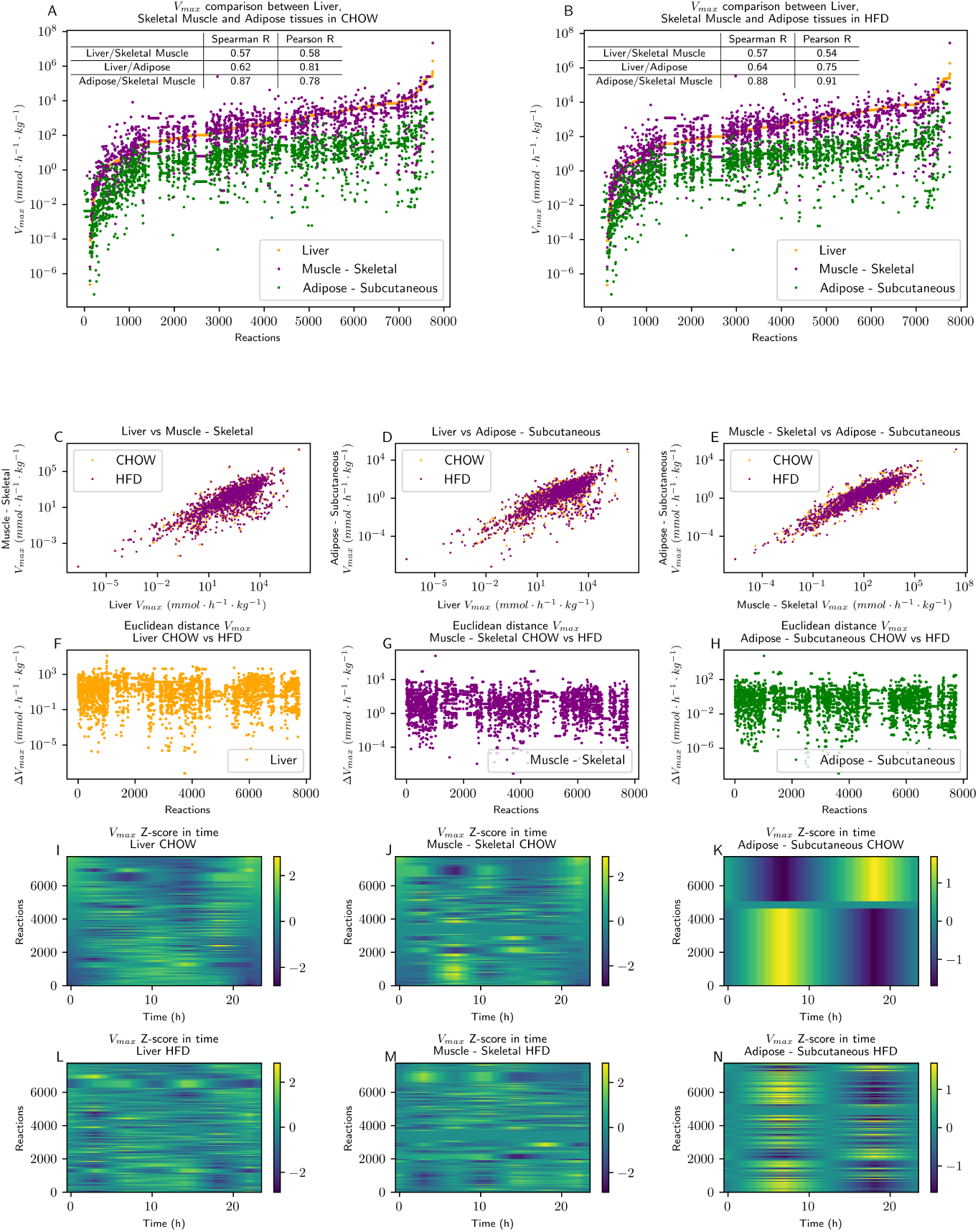
Characterization of the three tissues. In all the panels, the flux values represented are normalized over tissue mass to make them more comparable. In panels A and B, there is a overall representation of the maximum flux value for the three tissues and two dietary conditions at ZT 7. In both A and B the values have been sorted on the liver and it is possible to see that the skeletal muscle and the white adipose are very different from the liver while they share some similarities between themselves. In panels C to E we show the similarity between the *V*_*max*_ values by diet and tissue in a more direct comparison. In panels G to H, there is the intra-tissue distance between the two different diets. Finally, in panels I to N, there is the in-time representation of all six tissue and diet conditions extracted from the datasets. The values in these final panels have been normalized by reaction with a Z-score to more easily emphasize the in-time changes for each reaction. By any means, this representation is not interpretable as having 8000 circadian reactions with significative scores.

#### 4.5.2. Metabolomics

To validate the performances of the simulation being described, it is required to have a golden standard of metabolism in time. Unfortunately data on human are scarce and impossible to obtain, they also are subject to greater variability and usually restricted to bio-fluids, such as blood or urine (Dallmann et al., 2012).

Therefore we leveraged the availability of a very large dataset of murine metabolomics circadian data: The Atlas of Circadian Metabolism (Dyar et al., 2018). From the atlas we processed the data relative to liver, skeletal muscle, white adipose tissue and serum for both CHOW and HFD. Metabolomics data are produced extracting from the tissue a water soluble fraction and an insoluble fraction. These respectively contain the small polar molecules and the fats and other apolar molecules. Due to this extraction process, it is not possible to compare the relative abundance of these molecules to each other, but only between conditions.

We then map the metabolites present in the data to the model (detailed explanation in the upcoming sections). These data have many missing values that we decided to fill using a K-nearest-neighbor imputer from the python package scikit-learn (Pedregosa et al., 2011). We normalized the data with the geometric mean and interpolated them with a periodic cubic spline passing by the 6 or 7 averaged time-points available, like for the gene expression, therefore obtaining a function that we can interrogate using the time coordinate. The data available are reported in table 4.

**Table 4:** Data availability.

| Tissue | Condition | Data type | Time points | Source |
| --- | --- | --- | --- | --- |
| Liver | CHOW | Gene Expression | 7 | (Eckel-Mahan et al., 2013) |
|  |  | Metabolomics | 6 | (Dyar et al., 2018) |
|  | HFD | Gene Expression | 7 | (Eckel-Mahan et al., 2013) |
|  |  | Metabolomics | 6 | (Dyar et al., 2018) |
|  | Bmal WT | Gene Expression | 6 | (Koronowski et al., 2019; Smith et al., 2023) |
|  |  | Metabolomics | 6 | (Dyar et al., 2018) |
|  | Bmal KO | Gene Expression | 6 | (Koronowski et al., 2019; Smith et al., 2023) |
|  | Liver RE | Gene Expression | 6 | (Koronowski et al., 2019; Smith et al., 2023) |
|  | Muscle RE | Gene Expression | 6 | (Koronowski et al., 2019; Smith et al., 2023) |
|  | Double RE | Gene Expression | 6 | (Koronowski et al., 2019; Smith et al., 2023) |
| Skeletal Muscle | CHOW | Gene Expression | 6 | In house |
|  |  | Metabolomics | 6 | (Dyar et al., 2018) |
|  | HFD | Gene Expression | 6 | In house |
|  |  | Metabolomics | 6 | (Dyar et al., 2018) |
|  | Bmal WT | Gene Expression | 6 | (Koronowski et al., 2019; Smith et al., 2023) |
|  |  | Metabolomics | 6 | (Dyar et al., 2018) |
|  | Bmal KO | Gene Expression | 6 | (Koronowski et al., 2019; Smith et al., 2023) |
|  | Liver RE | Gene Expression | 6 | (Koronowski et al., 2019; Smith et al., 2023) |
|  | Muscle RE | Gene Expression | 6 | (Koronowski et al., 2019; Smith et al., 2023) |
|  | Double RE | Gene Expression | 6 | (Koronowski et al., 2019; Smith et al., 2023) |
| White adipose tissue | CHOW | Gene Expression | 2 (ZT0-12) | (Xin et al., 2022) |
|  |  | Metabolomics | 7 | (Dyar et al., 2018) |
|  | HFD | Gene Expression | 2 (ZT0-12) | (Xin et al., 2022) |
|  |  | Metabolomics | 6 | (Dyar et al., 2018) |

### 4.6. Diets and feeding

The model framework is able to accept any kind of feeding, provided that the molecules in the diet are present in the bloodstream. We took as standard diets the CHOW (Prolab^®^ RMH 2500 5P14*) and the HFD (Research Diets inc.^®^ D12492) from the atlas of circadian metabolism (Dyar et al., 2018). These two diets are representative of a balanced diet and a western world high fat diet for mice. We manually mapped the molecules present in the formulations to the model and scaled the caloric amount of the diets to 2500 *kcal*, sufficient to satisfy the caloric intake of an adult male. We also produced a high fat-high calories version of the HFD with 3500 *kcal*.

The feeding approach to the model is “intravenous” at scheduled timings. Although models exist for the absorption rate of the molecules in the diet via the intestines (Martins Conde et al., 2021), they are usually very limited both in data availability, number and kind of molecules studied. Therefore, to model food intake, we opted to create an *ad-hoc* equation which tries to reasonably approximate a physiological behavior (equation (1)). We assume that the food (for humans) is consumed at regular intervals, namely breakfast, lunch, and dinner respectively at 7:00, 13:00, and 20:00. We also assume that the food is processed by the digestive tract in 5 hours and the absorption is not uniform but peaks shortly after the beginning of the intake and slowly goes down to 0 in 5 hours. We then split the caloric intake into 1/5 for breakfast and 2/5 for lunch and dinner. We use the integral form of the equation evaluated between the time-step interval for the actual computation.

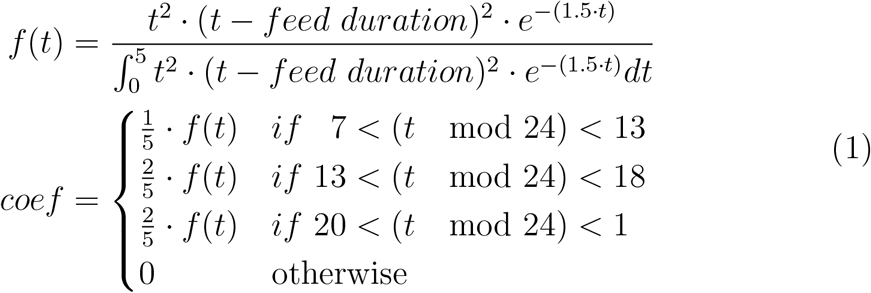

The equation (1) takes in input the time at which we want to get the amount of food absorbed. It is easy to integrate and *feed duration* is a parameter, in our model, set to 5 but it can be changed if needed. This equation allows for fast computation of the coefficients and is mathematically smooth and well behaved, as shown in figure 8. In particular, the equation has a first derivative equal to 0 at the time-points 0 and 5 from both sides, allowing a smooth transition from a feeding regimen to no intake. The coefficient is the multiplicative factor to the amount of metabolites added to the blood-stream. The actual multiplicative coefficient is always calculated analytically with the integral 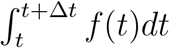 where *dt* is the time-step.

**Figure 8.**
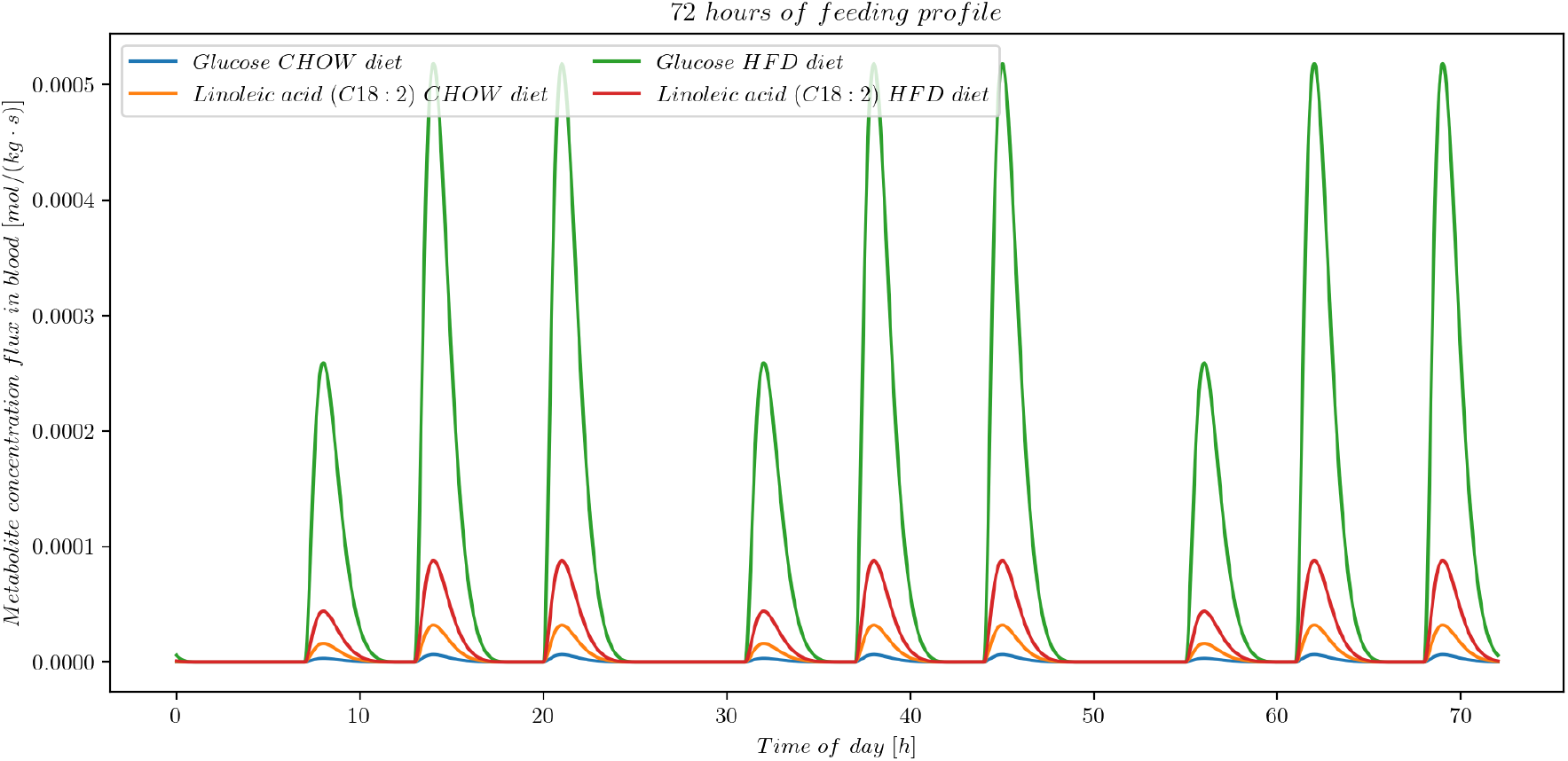
The profile of the food adsorption. Handmade to satisfy mathematical criteria (see equation (1)), it has a shape similar to the data derived from experiments (Martins Conde et al., 2021). In picture it is possible to see the difference between CHOW and HFD for two metabolites: glucose (CHOW in blue, HDF in green) and linoleic acid (CHOW in orange, HFD in red). The three periodic peaks represent breakfast, lunch, and dinner, and the function has a 24-hour period.

### 4.7. pH, ionic strength and osmotic balance

One of the biological systems main tasks is to keep pH and osmotic pressure balanced. This is achieved by buffer systems and ion and water channels and pumps. Our model considers two of the major buffer systems in humans, the bicarbonate and the phosphate buffers and keeps track of the osmotic balance between compartments.

#### 4.7.1. pH and ionic strength

After a wide search in literature, we found a series of methods to compute the pH of a buffered solution. The simplest, well-known, approach is the Henderson-Hasselbalch equation to calculate the pH of a single buffer solution. Unfortunately, this is not enough when the *pK*_*a*_s of the buffers are close to each other. Some resources like (Nguyen et al., 2009) (a paper widely criticized in literature) propose convoluted methods not well described and without an available code to apply. Some other approaches like (Stewart, 1978, 1981, 1983; Kellum, 2005) use electroneutrality of the solution as constraint, which we cannot use in our case for the double reason that cell compartments are not electroneutral and we do not have track of all charges present in solution (in particular we are missing the charges present on the proteins and the cell membranes).

We then devised a new, simpler method to compute the pH of the cell compartments in multiple buffered solutions leveraging the vast amount of information available on the metabolites concentrations. Given that we have available the initial pH, the increment of [*H*^+^], the *pk*_*a*_ of the species, the total concentration of the dissolved species ([*A*^−^] + [*AH*]) and the ionic strength of the solution, we use the following equations (2)

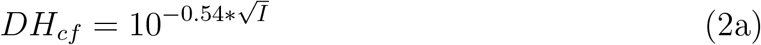

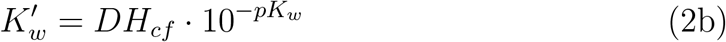

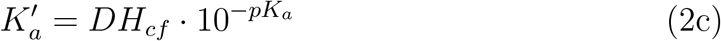

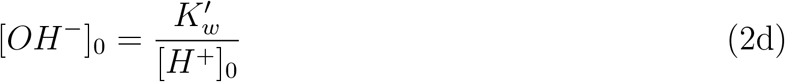

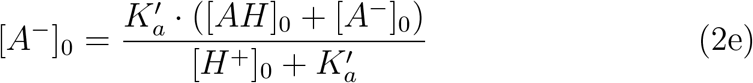

Here ([*AH*] + [*A*^−^]) is constant, *DH*_*cf*_ is the correction factor due to the Debye–Hückel theory (Holtzer, 1954) in water at 37^°^*C* and equations (2)d-e are the traditional equations to compute a dissociation equilibrium. At this point it is possible to add as many (monoprotic) acid or bases all following the same formulas 2c-e as one wishes. A key concept at this point is that all the increment of [*H*^+^] must be parted between the different equilibria, therefore the following equation (3) holds:

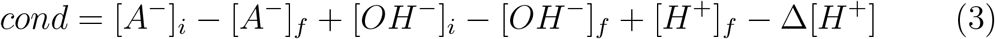

Where *i* means initial and *f* means final. We assert that the value of *cond* has to be 0 at the equilibrium since it means that all the Δ[*H*^+^] has been buffered, either by the buffer species or by the *OH*^−^ ions (changing the pH). The information regarding the initial pH is implicit in [*A*^−^]_*i*_ and [*OH*^−^]_*i*_. At this point we set the boundaries to the available values of [*H*^+^]_*f*_ to 10^−14^ and 1, we iteratively update the values of the final terms using equations (2), starting a bisection method with initial value for [*H*^+^]_*f*_ of 1. If *cond* ≥ 0 we change the upper boundary to [*H*^+^]_*f*_, otherwise we set the lower boundary to [*H*^+^]_*f*_. We then probe for the middle point until we get a convergence for *cond* to zero.

This method works for monoprotic acid (bases) and it suffices for our intents and purposes, since at physiological pH the phosphate and the carbonate buffers practically have only two microspecies present each. A microspecies is defined as follow: since each molecule containing an hydrogen atom bound to a highly electronegative atom is able to release it to water if the conditions are right, depending on the pH of the solution, the set of the protonated states of the molecule is called microspecies.

To estimate the charge of each compartment and compute the ionic strength, we used the same approach of Equilibrator 3.0 (Beber et al., 2022) to determine the distribution of microspecies. The idea is to calculate the energy of dissociation of each molecule using their *pKa* from literature or calculated by the software suite Chemaxon (Chemaxon). The distribution of microspecies is then computed using an approach similar to the partition function, which weights the abundance of each microspecies with the exponential of their energy, equation (4). This allows us to have the abundance of the charged particles in solution and, multiplying the distribution by the charge value of each microspecie, we can obtain the total estimated charge of the compartment. This is an estimation, therefore we are aware that it is wrong since we do not take into account proteins which usually have many negative charges exposed to the solvent and phospholipids which contain negative charges as well, but it is the best possible obtainable with the information available.

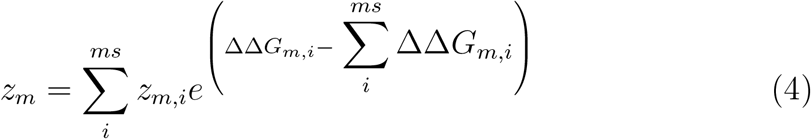

Where *ms* are the microspecies, ΔΔ*G*_*m,i*_ is the change of Gibbs free energy with respect to one of the microspecies set as reference and *z*_*m,i*_ is the charge of each microspecies, for example:

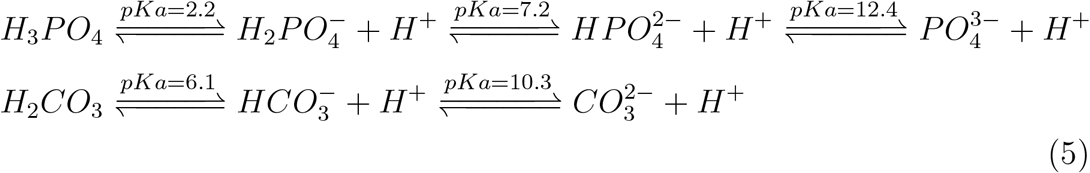

are the dissociation reactions of the two buffers and, consequently, the microspecies associated with the phosphoric acid and the carbonic acid. To be noted that the value of the second *pKa* for the phosphoric acid is equivalent to the *pH* of the cytosol and close to most of the compartments in the organism and similarly the first *pKa* of the carbonic acid, that is what makes this molecule one of the two most important buffers in the tissues.

As seen, the pH of the systems is also dependent on the ionic strength (IS) of the solution (for biological compartments, it is usually around 150 *mmol*/*kg*), which is, in turn, dependent on the charges of the species present in solution which are influenced by the IS and the pH in a circular fashion.

We calculate the IS as well with the following equation (6) where the concentrations *C*_*m*_ and the charges *z*_*m*_ are evaluated on the microspecies distribution

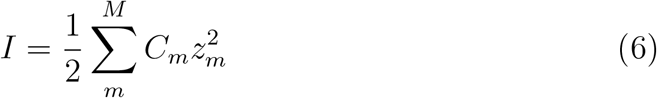

We, therefore, devised another iterative process to compute the same time pH, IS, and charges using also the equations from the Equilibrator (Beber et al., 2022) python packages. We did not go into detail to prove the convergence of the algorithm, but we tested it in several (also extreme) conditions, and, within the ranges allowed by our constraints, we observed that the results have always been consistent with realistic values. In line with Equilibrator 3.0 we used the Debye-Hückel equation to estimate the non-ideality of the activities in solution. Although we are aware that the equation is valid only considerably below the physiological IS. The implementation of the Pitzer equations for more concentrated solutions (Pitzer, 2018), and consequently also updating Equlibrator 3.0 to be in line with it, is beyond the scope of this work.

#### 4.7.2. Osmotic balance

The osmotic pressure is the sum of the concentration of all metabolites in solution (∑_*m*_ *c*_*m*_), where *c* is the concentration and *m* the small metabolites which we defined as having mass lower then 30kD, with *defined formula* (meaning it does not include unknown groups such as R, X and similar), excluding water. It is enforced to be around 300 mOsm/kg (Berman and Dugdale, 2023) in all compartments. It is kept in balance in the model constraining the reactions appropriately.

Moreover the two compartments of cytosol and nucleus are practically open from the point of view of a small molecule with size lower then 30 *kDa* through the opening of the nuclear pore (Paine et al., 1975; Alberts et al., 2002). Therefore the model considers the two compartments always in equilibrium within the 30 seconds long time-steps.

### 4.8 Solubility and water

The solubility of the metabolites in the system plays an important role for the organism and can determine even the insurgence of pathological diseases, such as kidney and gallbladder stones. In our modeling system we use the solubility to limit the concentration of metabolites, softly constraining them to have lower concentration then the computed solubility limits but, since the solubility predictor is still an approximation, allowing to overshoot these boundaries with a small penalty.

The vast majority of the metabolites do not have known solubility at 37°C, therefore we decided to use Chemaxon (Chemaxon) to compute the solubility of the species at the various compartments specific pH. We use the given estimation and clip the minimum and maximum value to a range of 10^−6^ *m* to 1 *m*. Practically, we do not have any metabolites with concentration higher than 165*mm* (*K*^+^ *mitochondrial*) and the overall sum of compartment metabolites concentration has to be lower than or equal to 300 mm (to remain under 300 mOsm/kg, see section 4.7). Although there are molecules with lower solubility than 10^−6^ *m*, we do not have accurate kinetic and spacial information and polar/apolar partitioning or availability of membranes which would allow for local increases of concentrations and activity of such metabolites that would justify smaller values in the modeling.

Water management in organisms is one of the most fundamental tasks performed by the systems. Therefore, in particular for land organisms, it has to be extremely tightly regulated. Due to our modeling choices we found extremely hard to keep track of water abundance in tissues and blood, adding on top bile secretion, urine excretion, the absence of a proper digestive and lung systems, sweating and a lymphatic system, it becomes impossible to proper manage water in the simulation. Moreover, most of the assumptions and laws underlying our model are true (more by necessity then by choice) only in the approximation of infinitely dilute solutions and the approximation does not hold in living systems. Also we do not keep track of the water that is normally adsorbed in large molecules and the colloidal nature of the fluids in the tissues. Finally, the changing water abundance would consequently change the concentration (and activity) of all the metabolites, and, although we keep track of the number of moles of the metabolites, we express concentration using molality, which is sensitive to water abundance. Changing the concentration to molar fraction might allow tracking and managing water in a more straightforward fashion but we did not explore the idea, also because we did not find any information regarding the connection between the species activity and the molar fraction in solutions.

### 4.9. Allosteric interactions

For optimal and correct behavior of the organism, the activity of the enzymes has to be tightly modulated at several levels distinguishable by the time scale and the energy involved in the regulation. The fastest regulatory mechanism for enzyme activity are the allosteric interactions with small molecules. Many pathways, for example glycolysis, are activated or repressed depending on the concentration of intermediate metabolites or metabolites representatives of the energy state of the cell. We, therefore, introduced an allosteric regulation that links metabolite concentrations and enzyme activity. We extracted the interactions from the allosteric database (Liu et al., 2020) and mapped the metabolites and the reactions against our model. Depending on the information reported on the database, we divided the metabolites as activators and inhibitors and modulated the activity of the linked reactions accordingly, using a sigmoid function with two parameters as shown in equation (7) and figure 9.

**Figure 9.**
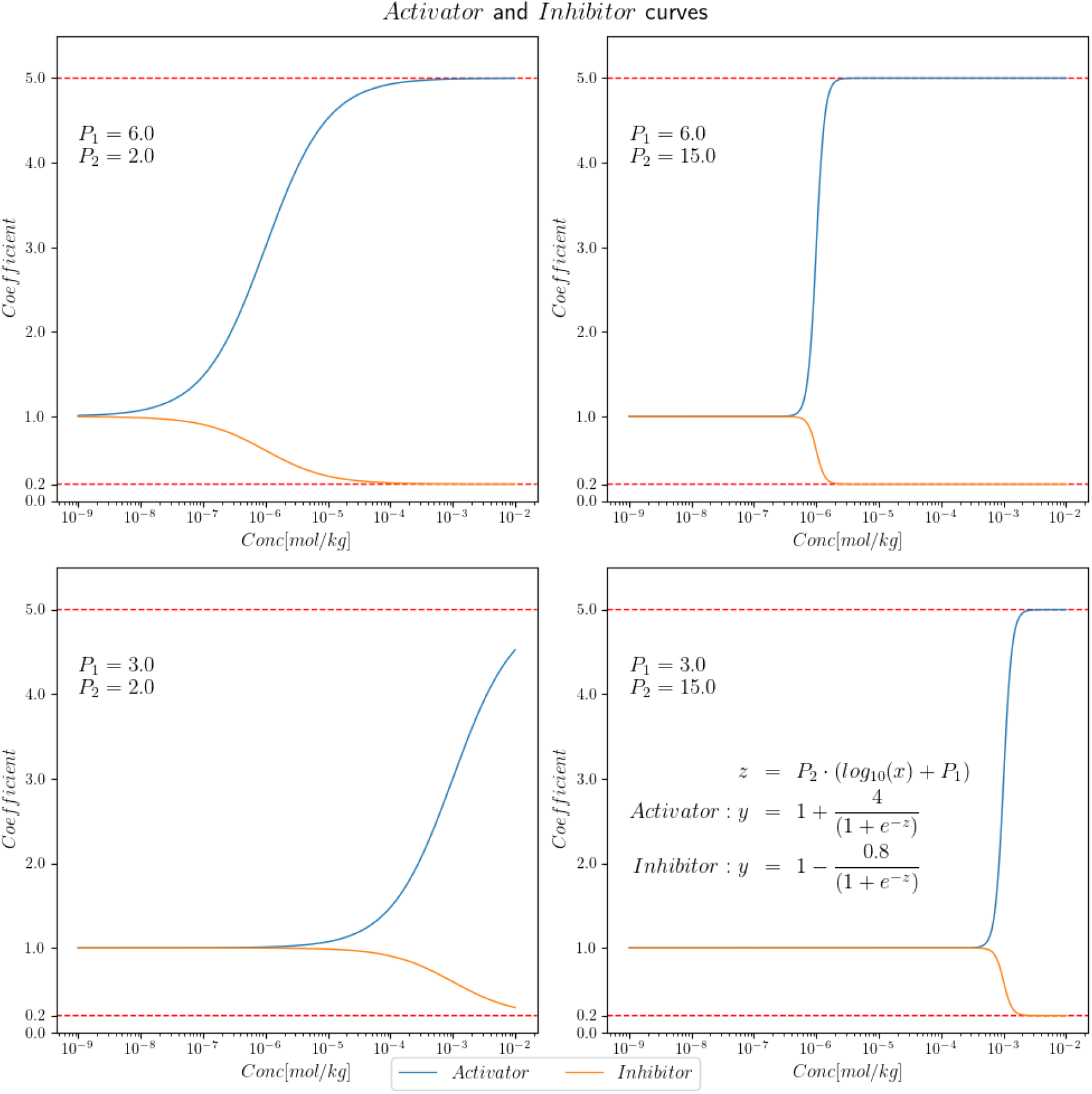
The sigmoid functions for the allosteric interaction of the metabolites with the relevant reactions. As we can see in the figure, the *Activator* and the *Inhibitor* coefficients are modeled by two different equations. The values of *P*_1_ and *P*_2_ parametrize the shape of the curve making the reaction more or less sensitive to the metabolite’s concentration (*x* axis) and shifting the concentration required to initiate the allosteric effect. The coefficient is multiplied by the reaction upper bound to limit the available flux. Notably, there is no possibility for a reaction to receive a 0 coefficient, and the multiplicative power of activation and inhibition cancel each other given the same parameters and concentrations.

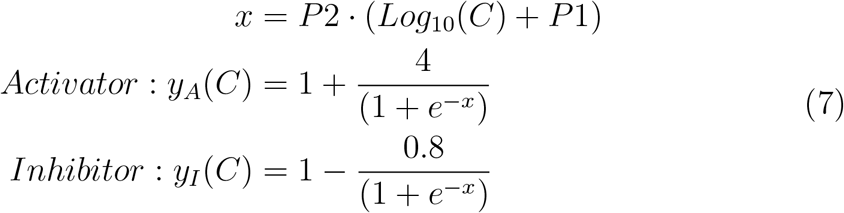

Where *C* is the concentration of the metabolites interacting with the enzyme. For simplicity, we will call the product of these coefficients *A*_*r*_

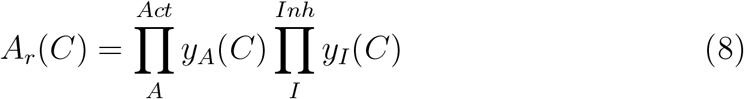

The coefficient *A*_*r*_ is always positive and, with the choice of 4 and 0.8 as numerators of for the logistic function, we ensure that they do not become too large or too small. The parameters *P* 1 and *P* 2 are unique for each relation and they are unknown. The method used to search for the appropriate values will be described later in the section.

We chose not to implement post-translational modification (e.g., phosphorylation) enzyme regulation at this iteration of the framework because it would bring a layer of complexity way above our scopes.

### 4.10. Metabolites

In this kind of models a metabolite is any object that participates in a reaction, most often it is a small molecule that can be represented with a structure and has some properties. Some metabolites, instead, are representing groups of metabolites, for example some fatty acids and molecules composed with them, which share the same sets of reactions, and these are called pool metabolites. This design choice has been present in metabolic networks for many years and it is quite laborious to revert, nevertheless we did our best to reduce the pools as much as possible to obtain the reactions related to the single molecules. Another kind of metabolite represents generic structures or substructures of large molecules and it is used to simulate specific reactions, for example phosphorylation of proteins (although without any additional effect then consuming the substrate) and DNA degradation.

A large effort has been put in defining the metabolites in the best possible way in order to maximize the matching with the external databases and reduce redundancy in the model.

#### 4.10.1. Standardization and matching

There are several ways to represent molecules computationally, it always depends on the degree of details required by the task. In our case, unfortunately, the level of detail required is very high. This is due to the extreme specificity of the enzymes to their substrate. One key example are the hexose sugars, fundamental molecules for life. These sugars, with brute formula *C*_6_*H*_12_*O*_6_, have a very large number of structures, conformation and chirality centers which all define the way they react and behave in the cell, therefore precautions have to be taken upon automatically define the structures of these molecules.

This brought the necessity of standardizing the representation of all molecules available. The main tool used for this task have been the Chemaxon suite (Chemaxon) and RDKit (RDKit) with a mix of automatic and manual approaches.

We also queried all relevant databases searching for molecules that were not reported with structural data, such as PubChem (Kim et al., 2023), Recon3D (Brunk et al., 2018), ChEBI (Hastings et al., 2016) and KEGG (Kanehisa and Goto, 2000). Where was not possible we wrote down the structure manually and inserted it in the database.

We also leveraged the database already present inside Equilibrator 3.0 (Beber et al., 2022), which is the software used to compute the properties of the molecules, in conjunction with Chemaxon, as explained in the upcoming section.

Once the metabolites in the model database have been standardized with the proper structure (primarily using SMILES format), they have been matched against the metabolomics data of mouse from the Atlas of Circadian Metabolism (Dyar et al., 2018) and the allosteric database (Liu et al., 2020). The automatic matching has not been very successful for the metabolomics data, therefore a mixed approach of string matching with the common names and manual curation has been deemed necessary. Against the allosteric database, instead, the information contained was almost exclusively structural and the database was not well organized in terms of connection, ids and names, therefore a complete manual curation had to be performed after the automatic matching.

This process emphasizes the problem of the past few decades of database matching with different entry names, the necessity of a standardization in the bioinformatics and cheminformatics community to achieve a unified way to represent items without risking overlapping or excessive generalization of terms.

#### 4.10.2. Properties calculations

Finally, to construct the model including thermodynamics and solubility information, it is required to calculate the 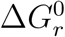 and the solubility of the molecules themselves. To do so the first thing required is the structural information of the metabolites, found and inserted in the model as explained in the previous section, then this information, in conjunction with compartment *pH*, ionic strength and electric potential can be submitted to the software Equilibrator 3.0 (Beber et al., 2022) to obtain the 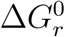. Equilibrator 3.0 uses a combination of machine learning, group contribution, component contribution and chemistry equations to obtain the desired value, it is also able to provide a confidence score for the prediction.

Equilibrator 3.0 leverages also the partition of metabolites in their microspecies, assumes diluted solution, includes the ionic strength with the Debye–Hückel equation and pH. Due to the complexity of the calculations and large number of reactions in the system, Equilibrator is too slow to be included directly in a dynamic simulation. Therefore we precomputed the values of the 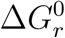 on an unevenly distributed grid of possible values for pH and IS, increasing the density of points where the gradient of the function becomes steep. In this way we constructed a linear interpolator from ℝ^2^ → ℝ for the single compartment reactions and from ℝ^4^ → ℝ for the multicompartment. The interpolated functions are much more lightweight to compute with respect to the calculations by the software and the error committed is minimal. For the values of electric potential across membranes and base pH values we used the default provided by Equilibrator in table 5.

**Table 5:** pH and electric potential of the compartments. The values of pH are adimensional, the values of electric potential, in *mV*, reported have the zero set to the cytosol.

| Compartment | pH | Electric potential<br>wrt Cytosol ($mV$) |
| --- | --- | --- |
| Cytosol | 7.2 | 0 |
| Mitochondria | 8 | -155 |
| Endoplasmatic<br>Reticulum | 7.2 | 0 |
| Nucleus | 7.2 | 0 |
| Peroxisome | 7.2 | 12 |
| Lysosome | 5.0 | 19 |
| Golgi<br>apparatus | 6.35 | 0 |
| Lipid Droplet/<br>Liposome | 7.2 | 0 |
| External Space | 7.4 | 18 |
| Blood | 7.4 | 30 |
| Red Blood<br>Cell Cytosol | 7.1 | 18 |
| Bile | 8.2 | 30 |
| Urine | 7.1 | 0 |

To estimate the 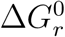 of the reaction for which Equilibrator was unable to compute a meaningful value (very high reported error), we decided to formulate an optimization problem in the form of a genetic algorithm, explained in the upcoming section, which determines all the free and unknown parameters of the simulation.

### 4.11. Metabolic Networks

Metabolic networks are a large collection of knowledge about metabolites and biochemical reactions. They set the relationships between such metabolites, how are they produced and consumed and in which proportion, and the reactions, which can be catalyzed by enzymes or spontaneous. The most advanced metabolic networks are called Genome-Scale Metabolic Models (GEMs) which encompass all possible (known) reactions inside a cell and transport between the different compartments.

Typically, GEMs are automatically constructed from other knowledge bases (Thiele and Palsson, 2010) and are unspecific, therefore, to obtain smaller and more targeted networks, they are routinely reduced in size, using information such as gene expression, representative of a specific cell type or tissue. A summary of the characteristics of some of the most popular and advanced metabolic networks is reported in table 6. As stated in the introduction, our starting metabolic network was chosen to be HGEM1.19.

**Table 6:**
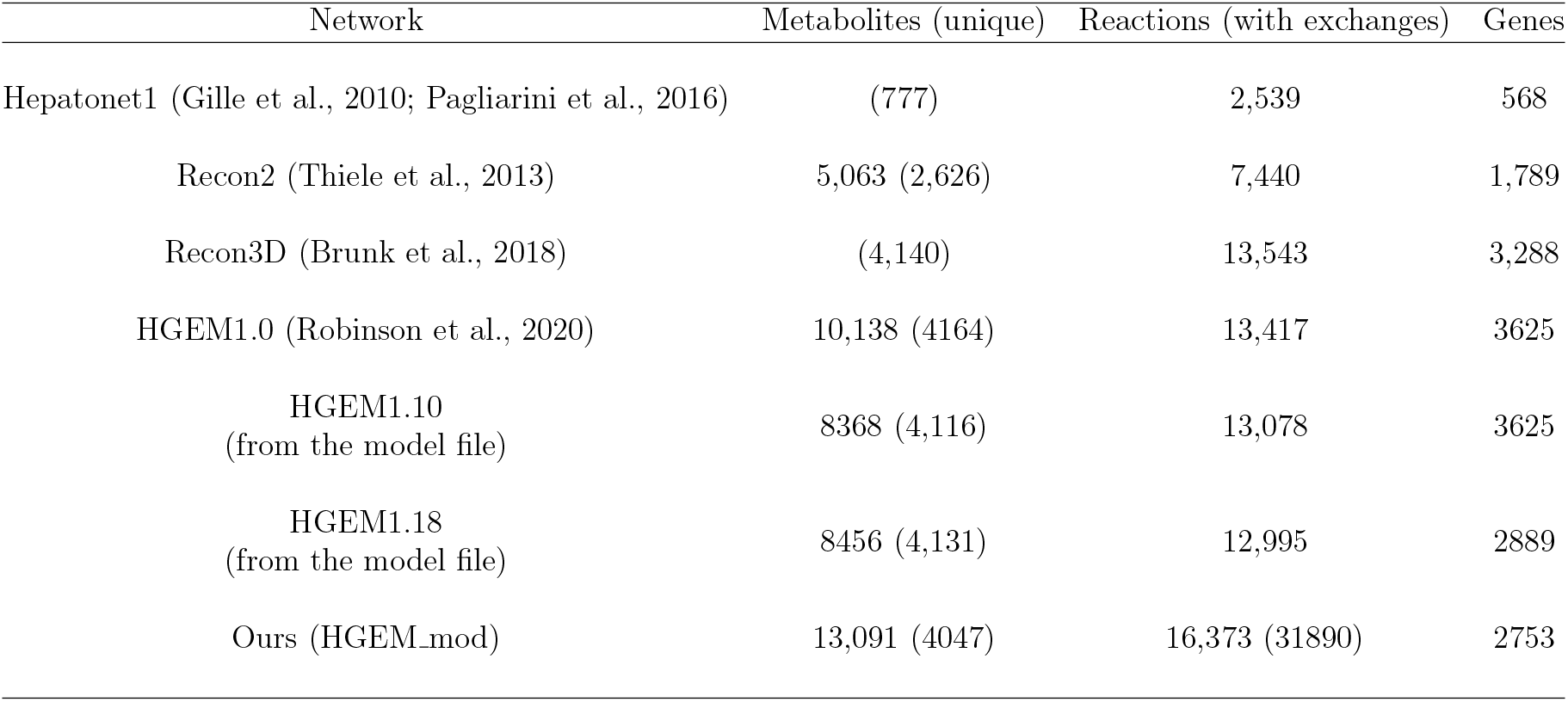
Characteristics of the different GEMs in literature.

### 4.12. Updates to HGEM1.19

Our workflow leverages COBRApy (the Python version of openCOBRA) (Heirendt et al., 2019) to read, modify and write HGEM1. COBRApy is extremely versatile and powerful in terms of data manipulation regarding GEMs, unfortunately, trying with our best efforts, we were unable to utilize it to our purposes being it extremely slow to update the model structure during a dynamic simulation. Therefore we devised a hybrid strategy that involves a preprocessing step of the model using the toolbox and the refinement of the problem definition and the optimization itself performed on a text based custom approach that allows us to have a speedup of many orders of magnitude.

In this section we will explain all the updates made to HGEM1.19 to get HGEM1.19+ and in the upcoming sections we will describe the algorithm we devised and the methods that we used to maximize computational efficiency.

#### 4.12.1. Pruning

The first thing we did was to remove the metabolite “glucose”. As explained in section 4.10.1, this key metabolite is very problematic from the identification and matching standpoint, therefore we collapsed it into the other, already present, metabolite *beta-D-glucose* and harmonized all reactions in this regard.

With a heuristic, we identified duplicated reactions and metabolites and removed them or collapsed them in one in the same way. We manually identified some problematic metabolites and reactions, either ill-defined or plain wrong, and removed them. Finally, with another heuristic loosely inspired by the Kirchhoff’s circuit laws, we removed isolated reactions and metabolites and dead ends.

We also removed all “BIOMASS” and similar reactions, since they are often tissue specific and are usually just sink reaction removing material from the system with very rigid proportions without properly dumping them in the appropriate location (such as the external environment).

#### 4.12.2. Nucleic acids

We added several pathways to the model. First of all we created a pathway to deal with DNA damage and degradation in the nucleus, in order to create an usable reaction to demand energy from the system with physiological values. In normal conditions, the DNA of a cell is continuously damaged and replaced. This happens mainly by the reaction with reactive oxygen species formed via chemical reactions or environmental ionizing radiation. From Chatterjee and Walker (2017) we extracted the average number of DNA bases damaged per hour per nucleus (total of around 3,000, which we multiplied by 10 to better estimate the energy consumption of the repair mechanism) and multiplied it with the number of nuclei in the tissue obtained in section 4.2. This gives us an energy consuming reaction which is enforced to have an estimated flux, set as constraint in the model.

We also moved RNA synthesis to the nucleus instead of the cytosol and added the exportin mechanism, consuming GTP, to move it outside.

#### 4.12.3. Bile

We added back the bile compartment and, consequently, changed all reactions of this excretion system from releasing the metabolites in the external space to the new compartment, from Boyer (2013) and Khonsary (2017) we extracted the physiological concentrations of the bile salts and molecules present in the compartment and added the ones missing from the model with the relevant transport reactions.

#### 4.12.4. Ions

The key actors in keeping the osmotic balance in the body are the small ions, specifically *Mg*^2+^ (Romani, 2011), *Na*^+^ (Xiong and Zhu, 2016), *K*^+^ (Xiong and Zhu, 2016), *Cl*^−^ (Jahn et al., 2015; Xiong and Zhu, 2016) and *Ca*^2+^ (Kellokumpu, 2019). These where fixed through the model, adding relevant transport reaction (either diffusive or active) that were missing. Keeping the homeostasis of ions is a big energy expenditure for the cells and it is enforced indirectly constraining the model to keep the osmotic balance in check.

#### 4.12.5. Blood and red blood cell

We added a blood and red blood cells compartments, as explained in section 4.3, the blood starts as a copy of the external space in terms of metabolites present and concentrations, excluding very large molecules such as heparan-sulfates, which are part of the intercellular matrix. The red blood cell sub-network is taken from Bordbar et al. (2011), with metabolites properly matched with the main network. This sub-network can only communicate with transport reactions with the blood, it also hosts the hemoglobin and the carbonic anhydrase reactions for gas transport.

#### 4.12.6. Glycogen and lipid droplet

We created a metabolite *glycogen storage* where the glycogen formed by the model can be stored and can be released with the proper stoichiometry in terms of glycogen/glucose ratio (30,000/1). We also removed the pool metabolite lipid droplet, instead created a compartment. In this compartments there can be stored all metabolites that were previously accumulated in the lipid droplet, it is also the largest compartment of the white adipose tissue and it occupies a considerable portion of the liver and the skeletal muscle.

#### 4.12.7. Other additions and modifications

We changed protein degradation, partially moving it inside the lysosomes; we added the myosin ATP consumption reaction with the proper rate (Johnson et al., 2019) for the skeletal muscle; we fixed the synthesis and degradation of dinucleosides oligophosphates (Lee et al., 2004); we added the glutamyl cycle (Kwiecień et al., 2007); we added DNA modification reaction DNA-5mC; we fixed RNA degradation; we removed *NH*_3_ as the species is not the most abundant microspecies between *NH*_3_ and 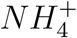 at cytosolic pH; we added transport reactions for the dihydroxiacetone phospate from the external space; we added the vitamin K and its related transport reactions and the vitamin K cycle; we fixed vitamin B12 and its related reactions; we added the histone modification reactions mediate by sirtuins being them an important connection between epigenetics and metabolism (Boon et al., 2020); we added the urine compartment and related transport reactions, as explained in section 4.3; we added part of the Kreb’s cycle in the nucleus (Kafkia et al., 2022); we added glucoronide, glucorosides and GSH adducts (mercapturic acids) synthesis pathways for all relevant molecules and transport to either bile or urine; finally, for the reasons stated in section 4.7, we added all small molecules from the cytosol to the nucleus and the relevant free diffusion reactions (Paine et al., 1975; Alberts et al., 2002).

### 4.13. A Thermo-Dyinamic Flux Balance Analysis

In this section we are going to build up all the components described previously to end up with the problem definition for our approach that we will call tdFBA.

Starting from equation (9):

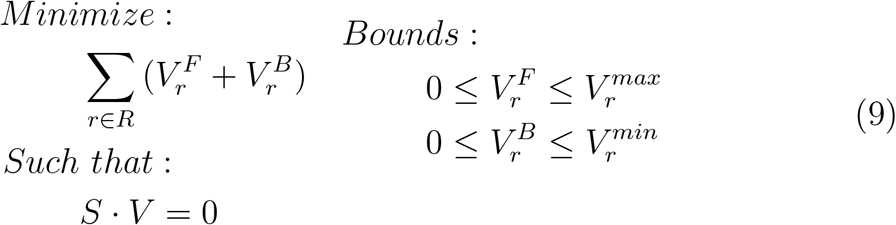

which defines a basic FBA optimization, we proceed to develop the dynamic model with a Static Optimization Approach (SOA) (Mahadevan et al., 2002). This means that, in general, we calculate at each time-step the steady state solution of the system and we carry over information to the next time-step with the relevant variables.

### 4.14. Introduction of time in the model

For each metabolite in each compartment we introduce in the model a set of reactions in the form:

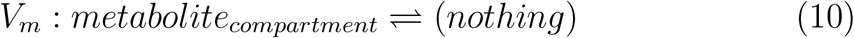

These reactions will enter in the stoichiometric matrix as constraints with an associated flux. These are sink/source reactions and they allow the model to remove or insert mass in the system and, with the appropriate bounds on their fluxes, they are treated as the linear approximation of the derivative in time of the concentration of the metabolites 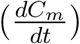. As with the other fluxes they are split in forward and backward reactions, with the former removing mass from the fluxes, therefore increasing concentration, and the latter adding mass to the fluxes, reducing the concentration.

At each time step the algorithm is required to solve a linear optimization problem that will update the concentrations according to the values computed for these derivatives, therefore, they have to be constrained according to the physiological conditions. Most importantly the concentrations must always be strictly positive otherwise they would break the logarithm operation that we take to compute the Δ*G*_*r*_ and softly bound by solubility. To enforce these constraints at each time step we introduced in the optimization problem also the length of the time step itself, in this way the concentrations can be computed during optimization and enforced to be within the established limits.

Therefore, for each metabolite are associated two linear constraints and two boundary conditions on the derivatives as it follows in equation (11):

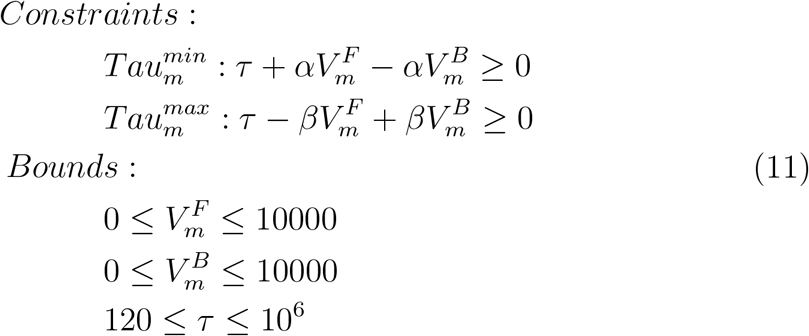

In the equation 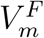 and 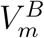 are the forward and backward variables representing the derivatives of the concentrations, while *τ* is the inverse of the time step size, spanning from 1*µs* to 30*s*. to The non zero and solubility constraints are taken into account in the values of *α* and *β*.

An exception has to be made for four compartments of the system, namely bile, blood, red blood cells and urine. These compartments are open therefore they have actual exchange reactions associated to them and, although they have the same reaction structure, are treated differently with respect to the *V*_*m*_.

#### 4.14.1. Urine

Starting with the simplest of the four, urine is a dummy compartment, it has no real role except hosting the exchange reactions. The concentration values of its metabolites are simply set to a very small number in order to ensure an always negative Δ*G*_*r*_ for the transport reactions from blood. The actual flux of these metabolites is set based on the concentration of the metabolites themselves in the blood, taking into account the values gathered from literature (section 4.2.4).

#### 4.14.2. Blood and red blood cells

Blood and red blood cells (in this subsection blood will mean both of the compartments unless specified) are modeled in a different way from the rest of the compartments, they do not actually have a “size” but they have a flow (of 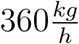, section 4.2). The values of their exchange reactions determine the concentration of the metabolites presented at each time-step to the tissues and therefore the amount available. For simplicity we assumed that the blood is completely recirculated every 60*s* (two full time-steps), this allows us to virtually split it into a portion in contact with the tissues and one “in the lungs” or in the rest of the circulatory system.

The part of the blood not in the tissues is enriched with *O*_2_ up to saturation, it has its *CO*_2_ removed and, if it is the case, it is enriched with nutrients from the diet. The rest of the blood carries the amount of metabolites equivalent to 30*s* of flow, with which the tissues can interact. In this scenario there are practically two separated exchange reactions for each metabolite in the compartments, a inward one that is set as a constraint with fixed value derived from the concentration of metabolites present in the “lungs” fraction of the blood, determined at the precedent time-step, and an outward one that has lower bound greater then zero and upper bound lower then the solubility of the metabolite. This enforces a constant flow of mass in the compartments. Meanwhile in the red blood cells, the reactions can happen normally, like in a standard FBA.

In case the time-step is lower then 30*s* we decided to have the model reach the exact 30*s* mark for the upcoming time-step in order to have the same blood flow for the entire span, dramatically simplifying the managing of the compartment.

The blood exiting the tissues is then pooled together in the “hearth” and the concentrations of the metabolites averaged on the values of the flows from the tissues before being redistributed at the next time-step. Calling these inward and outward fluxes *V* ^*i*^ and *V* ^*o*^ respectively we obtain equations (12).

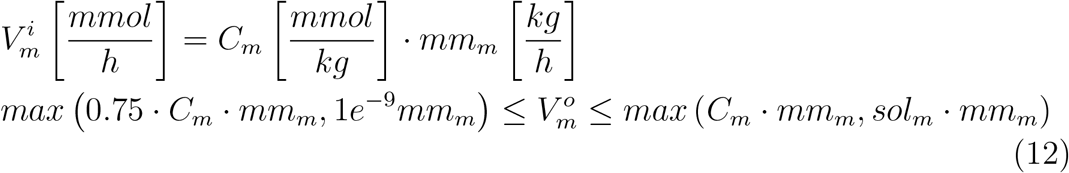

In this case *mm*_*m*_ is the proportion between the two compartments blood and red blood cell defined in table 2 multiplied by the mass flow of blood in the tissue (*F* ^*t*^) in 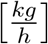. Given our choices of timings for the simulation and blood flow, the value already includes time in it in the right proportion.

#### 4.14.3. Bile

Bile shares some of the characteristics of blood, meaning that it has a flow (section 4.2) toward outside. There is no inward flow in the bile and the content is completely determined by the transport reactions from the cytosol of the liver, moreover, by design choice, we ignore water balance, see section 4.8.

In this case the concentration update is computed again using the the *V*_*m*_ of the compartment, and the exchange reaction acts only toward outside as a required sink. We define this behavior with equation (13).

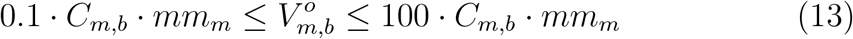

Where *C*_*m,b*_ is the concentration of the metabolite of the bile and *mm*_*m*_ is the mass of the liver times the proportion of bile as defined in table 2.

#### 4.14.4. Ions

A special mention has to be done for the major ion species in the system, *Ca*^2+^, *Mg*^2+^, *Na*^+^, *K*^+^ and *Cl*^−^. Their boundaries are more stringent in such a way that their concentrations cannot go under 0.8 and over 1.2 times the concentrations found in literature Wu et al. (2009); Brini and Carafoli (2011); Seifert et al. (2015); Jahn et al. (2015); Xu et al. (2016); Xiong and Zhu (2016); Kellokumpu (2019); Wang and Michalak (2020); Wu et al. (2023). A similar thought holds for the concentration of *H*^+^, although, with the buffer system in action, its variability is extremely low by default.

As a matter of fact, also the ions should be modeled with a buffering approach. *Mg*^2+^, for example, in physiological conditions, is always partially bound to the phosphate groups of nucleic acids, such as ADP and ATP, neutralizing them and removing it from the solution. This is a pivotal phenomenon for the management of the concentrations of both the nucleic acids and the magnesium itself in the mitochondria and it is related to availability of the substrates for the ATP-synthase. Although we do not model it explicitly, Equilibrator 3.0 takes the pMg (−*log*_10_[*Mg*^2+^]) into account when computing the Δ*G*_*r*_ of reactions involving nucleic acids.

#### 4.14.5. Concentration update

Finally we can define the concentration update equation for all metabolites as follows:

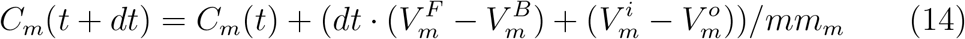

To be noted is that 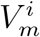 and 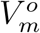 are not multiplied by time since in their *mm*_*m*_ it is already included. Moreover that all these operations assume vectorized form where values that are not relevant to each other are set to zero, therefore allowing a simple formulation of the equation instead of two sepa-rated equations (bile 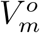 is also set to zero since the updating variables are the *V*_*m*_).

To update the concentration of blood and red blood cells we then take the values calculated from equation (14) and obtain the following:

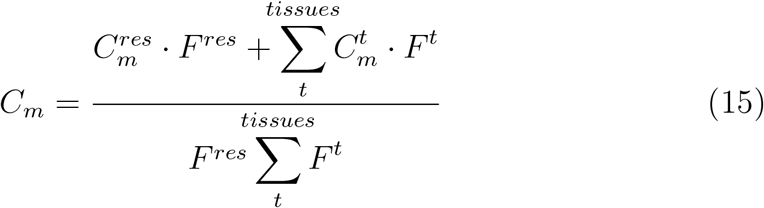

Where *F* ^*t*^ are the the flows in the tissues and *F* ^*res*^ is the residual flow available to the system which is supposed to irrorate the remaining tissues.

### 4.15. Introduction of thermodynamics in the model

One of the tasks for the development of this model is to introduce thermodynamics without the need of increasing the computational complexity of the optimization problem, like in Hoppe et al. (2007). The key factor is that this is a framework which includes time which is the main advantage of our approach, in fact we can encode in the model the equations to determine the directionality of the reactions outside the optimization.

Like for the tFBA, the same equations (16) and 17 still hold.

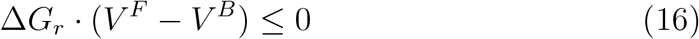

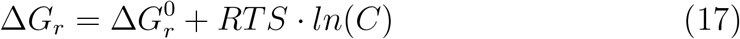

The main difference is that now the Δ*G*_*r*_ can be computed statically for each time-step with the concentrations available at the moment. Therefore we can plug in the values for the ionic strength and the *pH* calculated with the iterative method of section 4.7 on the precomputed values for the 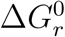 (section 4.10.2) and obtain the “constant” part. Then, with the concentrations available, we can apply equation (17).

The constraint of equation (16) can now be simplified by just setting to zero the values in the boundary section of the problem of the *V* ^*F*^, in case of positive Δ*G*_*r*_, or the *V* ^*B*^, in case of negative Δ*G*_*r*_. In case we obtain a value of Δ*G*_*r*_ of zero, or very close to, we allow flux in both directions for the time-step. Therefore, including also the allosteric coefficients obtained from equation (8), with the associated concentrations of the metabolites, we obtain the following structure for the boundaries of the fluxes, using the 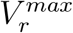 derived in section 4.5.1.

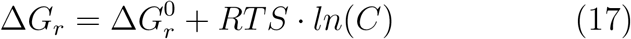

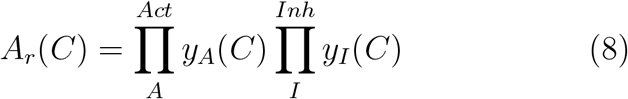

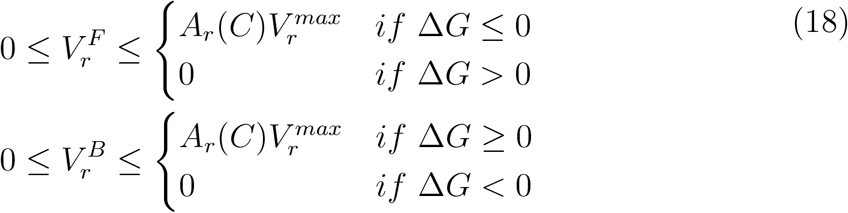

#### 4.15.1. Other constraints

With all the rest of the information gathered, we are able to construct the following additional constraints.

#### 4.15.2. Power

We introduce two constraints for the energy expended by the tissues, one for the upper limit and one for the lower. They have the the form of a sum of the multiplication of the Gibbs free energy by the flux of the associated reaction, equation (19). Calling the power for each tissue *W*_*t*_:

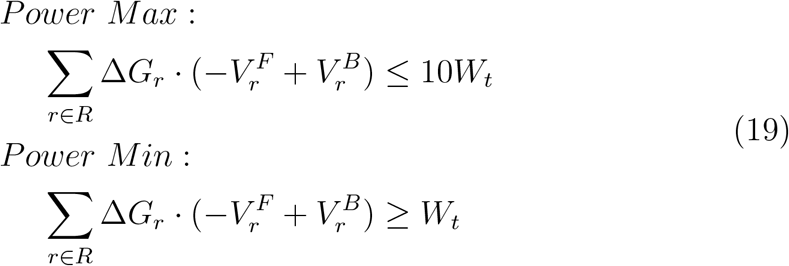

A minimal requirement of power implies that the energy in the system has to flow, this also limits, but does not eliminate, the possibility of futile loops inside the network.

#### 4.15.3. Osmotic balance

Since we do not have the concentrations inside the single optimization problems, to enforce the osmotic balance constraints we act on the derivatives of the concentrations, therefore, using the notation of section 4.14 we can build the following equation (20).

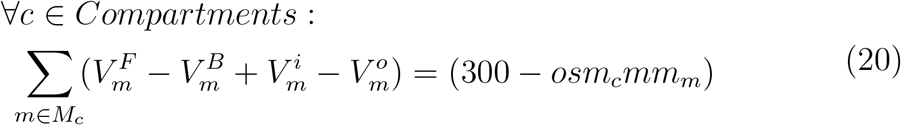

Where *osm*_*c*_ is the osmolarity of the compartment, of course, for blood and red blood cells, the values of 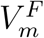 and 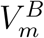 will be zero. Instead, for all the other compartments, 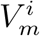 and 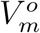 will be zero. This allows the system to return to the osmolarity of 300*mOsm* in case the constraint has to be relaxed to make the model feasible.

#### 4.15.4. Nucleus-cytoplasm transport

To enforce the fact that the two compartments are always in perfect equilibrium on the timescale of 30*s*, we add another constraint on the derivatives *V*_*m*_ associated to the two compartments, equation (21).

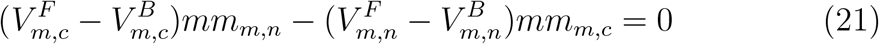

#### 4.15.5. Fat

Since we know the molecular mass of the metabolites present in the lipid droplet, we decided to enforce a minimal and maximal quantity of fat present in the tissues, based on the size and proportion of the compartment with respect to the entire tissue, equation (22).

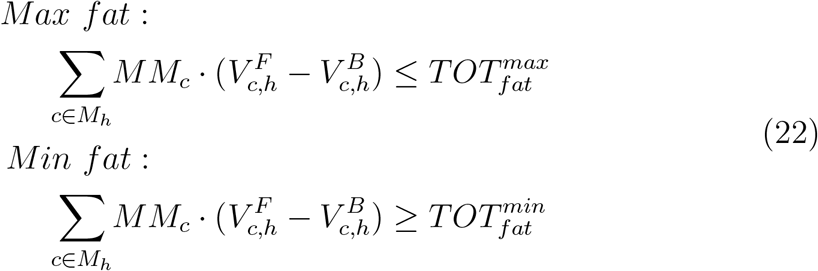

### 4.16. Objective function

Finally we have to define an objective function for the optimization problem. By design, we already excluded the biomass reactions from the model, since we are not interested in the growth of the organism.

We decided to test four different combinations of coefficients to assign to the flux values *V* in a maximization problem: zero, one, minus one, Δ*G*_*r*_ and the expression value before combining it with the *k*_*cat*_. For reactions without associated genes, instead, we opted to assign a negative coefficient to minimize them.

Moreover, we added to the objective function the maximization of the unbalanced transport reactions, the minimization of the derivatives of the metabolites, unless they reach values too small or too far from their initial concentrations, in which case we promote the opportune direction to return toward the previous values.

The objective function is then equation (23).

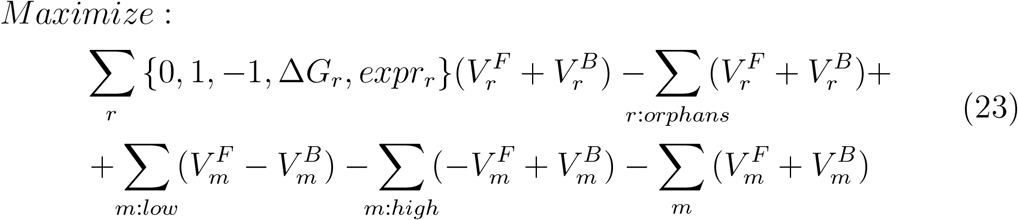

### 4.17. Final optimization problem

We can now assemble all the pieces together to formulate the optimization problem. Practically, this becomes a file of a few megabytes of text, which is then compiled like a form by the software with all the relevant numbers required. This solves the issue of constructing the problem with COBRA which is extremely time consuming, instead with this approach, we segregate most of this time in a preprocessing step of construction. Equation24 is the complete formulation.

Outside the optimization problems, there are all the updates to the states of the model as described by equations (14), (15) and the calculations for pH and ionic strength (section 4.7), Δ*G*_*r*_ (equation (17)) and allosteric coefficients (equation (8)). Each optimization problem is solved independently for each tissue with the IBM CPLEX optimizer. The framework allows to define the diet which will change the food intake of the model and the gene expression, accordingly. Moreover, it is possible to terminate the simulation by defining the number of steps to simulate or the total required time in hours. All the software has been optimized to maximize performances and run in parallel in order to be deployable on cluster and supercomputers.

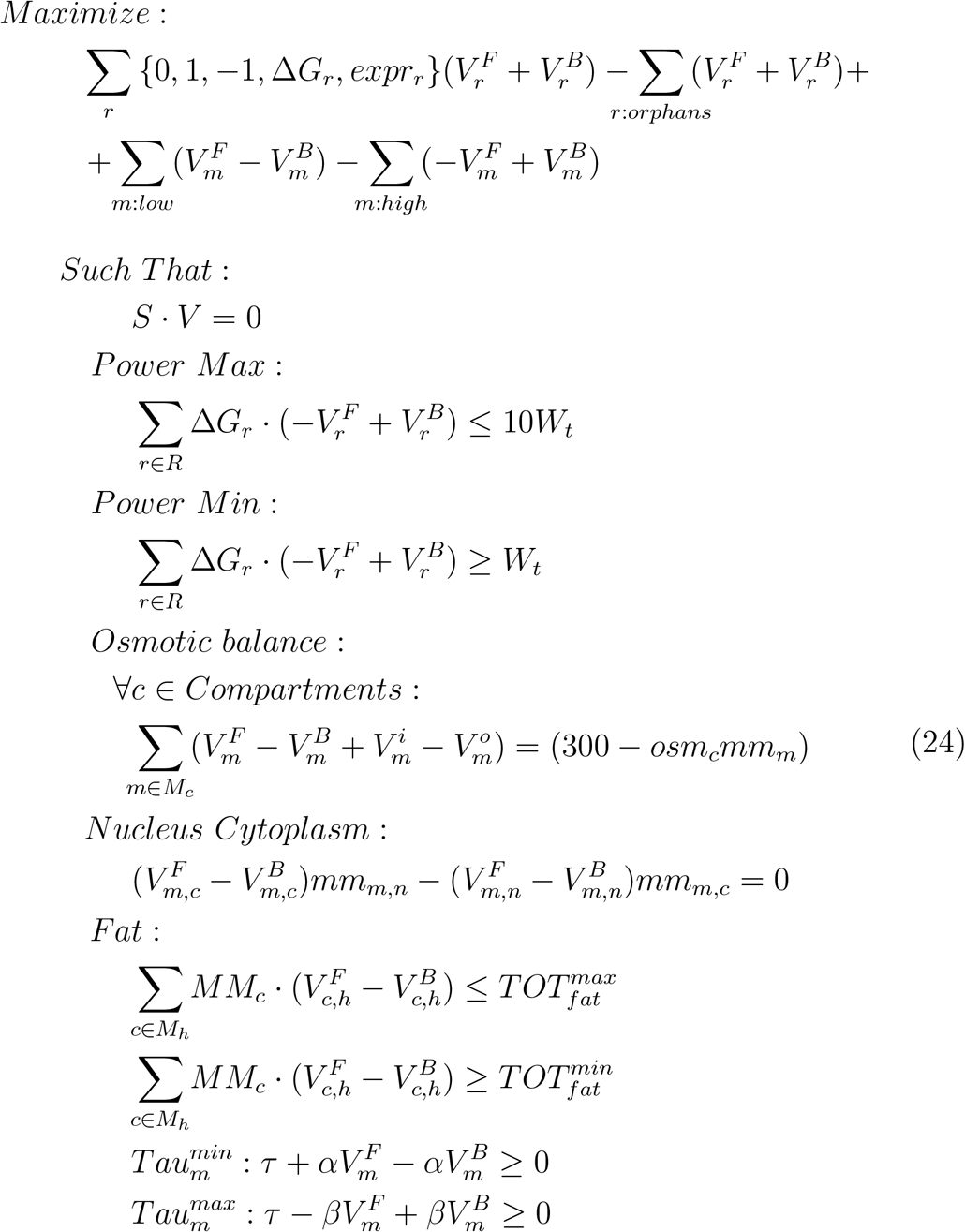

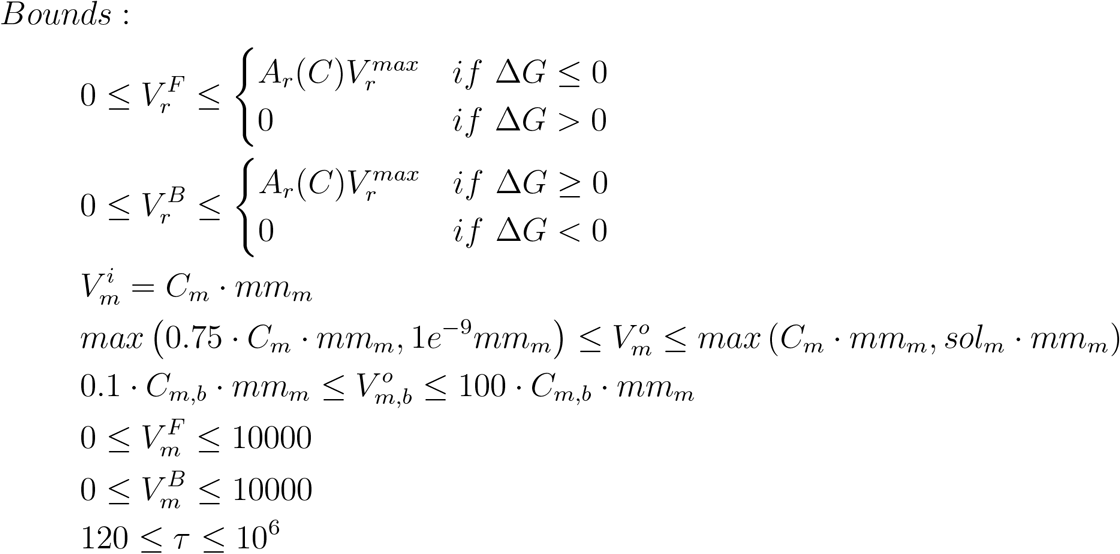

### 4.18. Missing parameters search

To define the coefficients for the fluxes part of the objective, the values for the 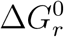 poorly defined by Equilibrator and the parameters for the allosteric coefficients, we decided to perform a search using a genetic algorithm. These optimization algorithms are among the most flexible available, with the drawback of not guaranteeing convergence, not even in local optima.

The genetic algorithm employed uses randomly selected values for the unknown parameters of the model as genes, runs the simulation for up to 48 hours, with stopping conditions in case the quality of the simulation degrades due to too many infeasibilities in the optimization problems. For example, if a Δ*G* becomes too large, it can force the model to completely deplete one or more metabolites. These metabolites may be substrates for other reactions or precursors for imposed demands. In this case, the model cannot be solved with the most stringent formulation, so the boundaries on the derivatives of the concentration of the metabolites are relaxed until a feasible solution is found. If too many relaxations occur in a row, one or more derivatives can become too large and the time-step has to be reduced accordingly. In this case if too many consecutive time-steps are less then 30*s*, the simulation is interrupted early.

The genetic algorithm algorithm runs a population of 100 simulations in parallel for each generation, then with a scoring function proportional to the time simulated and the correlation between the output of the model and the real murine metabolic circadian data for the CHOW diet, it selects the best fit members of the population, performs cross-over on the vector of the genes, mutations and, keeping the top ten members moves on to the next generation.

In this way, we estimated a set of parameters based only on one condition and selected the top 30 models across the generations to perform predictions on the other datasets of HFD and selective Bmal RE experiments, see table 4.

## References

Adkins, A.N., Dewald, J.P., Garmirian, L.P., Nelson, C.M., Murray, W.M., 2021. Serial sarcomere number is substantially decreased within the paretic biceps brachii in individuals with chronic hemiparetic stroke. Proceedings of the National Academy of Sciences of the United States of America 118. URL: https://pnas.org/doi/full/10.1073/pnas.2008597118, doi:10.1073/pnas.2008597118.

Alberts, B., Jhnson, A., Lewis, J., Raff, M., Roberts, K., Walter, P., 2002. The Transport of Molecules between the Nucleus and the Cytosol. volume 2154. URL: http://www.ncbi.nlm.nih.gov/books/NBK26932/figure/.

Asher, G., Sassone-Corsi, P., 2015. Time for food: The intimate interplay between nutrition, metabolism, and the circadian clock. Cell 161, 84–92. URL: https://linkinghub.elsevier.com/retrieve/pii/S0092867415003025, doi:10.1016/j.cell.2015.03.015.

Beber, M.E., Gollub, M.G., Mozaffari, D., Shebek, K.M., Flamholz, A.I., Milo, R., Noor, E., 2022. EQuilibrator 3.0: A database solution for thermodynamic constant estimation. Nucleic Acids Research 50, D603–D609. URL: https://doi.org/10.1093/nar/gkab1106, doi:10.1093/nar/gkab1106.

Berman, J., Dugdale, D.C., 2023. Osmolality - blood test. URL: https://www.mountsinai.org/health-library/tests/osmolality-blood-test.

Boon, R., Silveira, G.G., Mostoslavsky, R., 2020. Nuclear metabolism and the regulation of the epigenome. Nature Metabolism 2, 1190–1203. URL: https://www.nature.com/articles/s42255-020-00285-4, doi:10.1038/s42255-020-00285-4.

Bordbar, A., Jamshidi, N., Palsson, B.O., 2011. IAB-RBC-283: A proteomically derived knowledge-base of erythrocyte metabolism that can be used to simulate its physiological and patho-physiological states. BMC Systems Biology 5, 110. URL: https://bmcsystbiol.biomedcentral.com/articles/10.1186/1752-0509-5-110, doi:10.1186/1752-0509-5-110.

Boyer, J.L., 2013. Bile formation and secretion, in: Comprehensive Physiology. Wiley. volume 3, pp. 1035–1078. URL: https://onlinelibrary.wiley.com/doi/10.1002/cphy.c120027, doi:10.1002/cphy.c120027.

Brini, M., Carafoli, E., 2011. The Plasma Membrane Ca2+ ATPase and the plasma membrane Sodium Calcium exchanger cooperate in the regulation of cell Calcium. Cold Spring Harbor Perspectives in Biology 3, 1–15. URL: http://cshperspectives.cshlp.org/lookup/doi/10.1101/cshperspect.a004168, doi:10.1101/cshperspect.a004168.

Brunk, E., Sahoo, S., Zielinski, D.C., Altunkaya, A., Dräger, A., Mih, N., Gatto, F., Nilsson, A., Andres Preciat Gonzalez, G., Aurich, M.K., Prlić, A., Sastry, A., Danielsdottir, A.D., Heinken, A., Noronha, A., Rose, P.W., Burley, S.K., Fleming, R.M.T., Nielsen, J., Thiele, I., Palsson, B.O., Preciat Gonzalez, G.A., Aurich, M.K., Prlic, A., Sastry, A., Danielsdottir, A.D., Heinken, A., Noronha, A., Rose, P.W., Burley, S.K., Fleming, R.M.T., Nielsen, J., Thiele, I., Palsson, B.O., Prlić, A., Sastry, A., Danielsdottir, A.D., Heinken, A., Noronha, A., Rose, P.W., Burley, S.K., Fleming, R.M.T., Nielsen, J., Thiele, I., Palsson, B.O., 2018. Recon3D enables a three-dimensional view of gene variation in human metabolism. Nature Biotechnology 36, 272–281. URL: http://www.nature.com/articles/nbt.4072 http://dx.doi.org/10.1038/nbt.4072 http://vmh.life., doi:10.1038/nbt.4072.

Bryk, A.H., Wiśniewski, J.R., 2017. Quantitative Analysis of Human Red Blood Cell Proteome. Journal of Proteome Research 16, 2752–2761. URL: https://pubs.acs.org/sharingguidelines, doi:10.1021/acs.jproteome.7b00025.

Cameron, V.A., Autelitano, D.J., Evans, J.J., Ellmers, L.J., Espiner, E.A., Gary Nicholls, M., Mark Richards, A., 2002. Body-size dependence of resting energy expenditure can be attributed to nonenergetic homogeneity of fat-free mass. American Journal of Physiology - Endocrinology and Metabolism 282, E132–E138. URL: https://www.physiology.org/doi/10.1152/ajpendo.2002.282.1.E132, doi:10.1152/ajpendo.2002.282.1.e132.

Chang, A., Jeske, L., Ulbrich, S., Hofmann, J., Koblitz, J., Schomburg, I., Neumann-Schaal, M., Jahn, D., Schomburg, D., 2021. BRENDA, the ELIXIR core data resource in 2021: New developments and updates. Nucleic Acids Research 49, D498–D508. URL: https://academic.oup.com/nar/article/49/D1/D498/5992283, doi:10.1093/nar/gkaa1025.

Chatterjee, N., Walker, G.C., 2017. Mechanisms of DNA damage, repair, and mutagenesis. Environmental and Molecular Mutagenesis 58, 235–263. URL: https://onlinelibrary.wiley.com/doi/10.1002/em.22087, doi:10.1002/em.22087.

Chemaxon,. Calculators & predictors. URL: https://chemaxon.com/calculators-and-predictors.

Dadson, P., Ferrannini, E., Landini, L., Hannukainen, J.C., Kalliokoski, K.K., Vaittinen, M., Honka, H., Karlsson, H.K., Tuulari, J.J., Soinio, M., Salminen, P., Parkkola, R., Pihlajamäki, J., Iozzo, P., Nuutila, P., 2017. Fatty acid uptake and blood flow in adipose tissue compartments of morbidly obese subjects with or without type 2 diabetes: Effects of bariatric surgery. American Journal of Physiology - Endocrinology and Metabolism 313, E175–E182. URL: https://www.physiology.org/doi/10.1152/ajpendo.00044.2017, doi:10.1152/ajpendo.00044.2017.

Dallmann, R., Viola, A.U., Tarokh, L., Cajochen, C., Brown, S.A., 2012. The human circadian metabolome. Proceedings of the National Academy of Sciences of the United States of America 109, 2625–2629. URL: https://www.pnas.org/doi/abs/10.1073/pnas.1114410109, doi:10.1073/pnas.1114410109.

Dyar, K.A., Lutter, D., Artati, A., Ceglia, N.J., Liu, Y., Armenta, D., Jastroch, M., Schneider, S., de Mateo, S., Cervantes, M., Abbondante, S., Tognini, P., Orozco-Solis, R., Kinouchi, K., Wang, C., Swerdloff, R., Nadeef, S., Masri, S., Magistretti, P., Orlando, V., Borrelli, E., Uhlenhaut, N.H., Baldi, P., Adamski, J., Tschöp, M.H., Eckel-Mahan, K., Sassone-Corsi, P., 2018. Atlas of Circadian Metabolism Reveals System-wide Coordination and Communication between Clocks. Cell 174, 1571–1585.e11. doi:10.1016/j.cell.2018.08.042.

Eckel-Mahan, K.L., Patel, V.R., De Mateo, S., Orozco-Solis, R., Ceglia, N.J., Sahar, S., Dilag-Penilla, S.A., Dyar, K.A., Baldi, P., Sassone-Corsi, P., 2013. Reprogramming of the circadian clock by nutritional challenge. Cell 155, 1464–1478. URL: http://dx.doi.org/10.1016/j.cell.2013.11.034, doi:10.1016/j.cell.2013.11.034.

Elia, M., 1992. Organ and tissue contribution to metabolic rate. Energy metabolism. Tissue determinants and cellular corrolaries, 61–77.

Esteller, A., 2008. Physiology of bile secretion. World Journal of Gastroenterology 14, 5641–5649. URL: http://www.wjgnet.com/1007-9327/full/v14/i37/5641.htm, doi:10.3748/wjg.14.5641.

Gille, C., Bölling, C., Hoppe, A., Bulik, S., Hoffmann, S., Hübner, K., Karlstädt, A., Ganeshan, R., König, M., Rother, K., Weidlich, M., Behre, J., Holzhütter, H.G., 2010. HepatoNet1: a comprehensive metabolic reconstruction of the human hepatocyte for the analysis of liver physiology. Molecular systems biology 6, 411. URL: http://www.ncbi.nlm.nih.gov/pubmed/20823849 http://www.pubmedcentral.nih.gov/articlerender.fcgi?artid=PMC2964118, doi:10.1038/msb.2010.62.

Hall, J.E., Guyton, A.C., 2006. Guyton and hall textbook of medical physiology. 11 ed., Elsevier Inc.

Hastings, J., Owen, G., Dekker, A., Ennis, M., Kale, N., Muthukrishnan, V., Turner, S., Swainston, N., Mendes, P., Steinbeck, C., 2016. ChEBI in 2016: Improved services and an expanding collection of metabolites. Nucleic Acids Research 44, D1214–D1219. URL: https://academic.oup.com/nar/article-lookup/doi/10.1093/nar/gkv1031, doi:10.1093/nar/gkv1031.

Heirendt, L., Arreckx, S., Pfau, T., Mendoza, S.N., Richelle, A., Heinken, A., Haraldsdóttir, H.S., Wachowiak, J., Keating, S.M., Vlasov, V., Magnusdóttir, S., Ng, C.Y., Preciat, G., Žagare, A., Chan, S.H.J., Aurich, M.K., Clancy, C.M., Modamio, J., Sauls, J.T., Noronha, A., Bordbar, A., Cousins, B., El Assal, D.C., Valcarcel, L.V., Apaolaza, I., Ghaderi, S., Ahookhosh, M., Ben Guebila, M., Kostromins, A., Sompairac, N., Le, H.M., Ma, D., Sun, Y., Wang, L., Yurkovich, J.T., Oliveira, M.A.P., Vuong, P.T., El Assal, L.P., Kuperstein, I., Zinovyev, A., Hinton, H.S., Bryant, W.A., Aragón Artacho, F.J., Planes, F.J., Stalidzans, E., Maass, A., Vempala, S., Hucka, M., Saunders, M.A., Maranas, C.D., Lewis, N.E., Sauter, T., Palsson, B.Ø., Thiele, I., Fleming, R.M.T., 2019. Creation and analysis of biochemical constraint-based models using the COBRA Toolbox v.3.0. Nature Protocols 14, 639–702. URL: http://www.nature.com/articles/s41596-018-0098-2, doi:10.1038/s41596-018-0098-2.

Holtzer, A.M., 1954. The collected papers of Peter J. W. Debye. Interscience, New York-London, 1954. txxi + 700 pp., $9.50. Journal of Polymer Science 13, 548–548. URL: https://onlinelibrary.wiley.com/doi/10.1002/pol.1954.120137203, doi:10.1002/pol.1954.120137203.

Hoppe, A., Hoffmann, S., Holzhütter, H.G., 2007. Including metabolite concentrations into flux balance analysis: Thermodynamic realizability as a constraint on flux distributions in metabolic networks. BMC Systems Biology 1. doi:10.1186/1752-0509-1-23.

Hughes, M.E., Hogenesch, J.B., Kornacker, K., 2010. JTK-CYCLE: An efficient nonparametric algorithm for detecting rhythmic components in genome-scale data sets. Journal of Biological Rhythms 25, 372–380. URL: https://journals.sagepub.com/doi/10.1177/0748730410379711, doi:10.1177/0748730410379711.

Jahn, S.C., Rowland-Faux, L., Stacpoole, P.W., James, M.O., 2015. Chloride concentrations in human hepatic cytosol and mitochondria are a function of age. Biochemical and Biophysical Research Communications 459, 463–468. URL: https://linkinghub.elsevier.com/retrieve/pii/S0006291X15003757, doi:10.1016/j.bbrc.2015.02.128.

Jansson, P.A.E., Gudbjörnsdóttir, H.S., Andersson, O.K., Lönnroth, P.N., 1996. The effect of metformin on adipose tissue metabolism and peripheral blood flow in subjects with NIDDM. Diabetes Care 19, 160–164. URL: https://diabetesjournals.org/care/article/19/2/160/19776/The-Effect-of-Metformin-on-Adipose-Tissue, doi:10.2337/diacare.19.2.160.

Jiang, Y., Li, L., Chen, X., Liu, J., Yuan, J., Xie, Q., Han, H., 2021. Three-dimensional ATUM-SEM reconstruction and analysis of hepatic endoplasmic reticulum organelle interactions. Journal of Molecular Cell Biology 13, 636–645. URL: https://academic.oup.com/jmcb/article/13/9/636/6287621, doi:10.1093/jmcb/mjab032.

Johnson, C.A., Walklate, J., Svicevic, M., Mijailovich, S.M., Vera, C., Karabina, A., Leinwand, L.A., Geeves, M.A., 2019. The ATPase cycle of human muscle myosin II isoforms: Adaptation of a single mechanochemical cycle for different physiological roles. Journal of Biological Chemistry 294, 14267–14278. URL: https://linkinghub.elsevier.com/retrieve/pii/S0021925820349875, doi:10.1074/jbc.RA119.009825.

Kafkia, E., Andres-Pons, A., Ganter, K., Seiler, M., Smith, T.S., Andrejeva, A., Jouhten, P., Pereira, F., Franco, C., Kuroshchenkova, A., Leone, S., Sawarkar, R., Boston, R., Thaventhiran, J., Zaugg, J.B., Lilley, K.S., Lancrin, C., Beck, M., Patil, K.R., 2022. Operation of a TCA cycle subnetwork in the mammalian nucleus. Science Advances 8. URL: https://www.science.org/doi/10.1126/sciadv.abq5206, doi:10.1126/sciadv.abq5206.

Kanehisa, M., Goto, S., 2000. KEGG: Kyoto Encyclopedia of Genes and Genomes. Technical Report 1. URL: http://www.genome.ad.jp/kegg/.

Kellokumpu, S., 2019. Golgi pH, ion and redox homeostasis: How much do they really matter? Frontiers in Cell and Developmental Biology 7. URL: https://www.frontiersin.org/article/10.3389/fcell.2019.00093/full, doi:10.3389/fcell.2019.00093.

Kellum, J.A., 2005. Clinical review: Reunification of acid-base physiology. Critical Care 9, 500–507. doi:10.1186/cc3789.

Khonsary, S.A., 2017. Guyton and Hall: Textbook of Medical Physiology.. volume 8. doi:10.4103/sni.sni_327_17.

Kim, S., Chen, J., Cheng, T., Gindulyte, A., He, J., He, S., Li, Q., Shoemaker, B.A., Thiessen, P.A., Yu, B., Zaslavsky, L., Zhang, J., Bolton, E.E., 2023. PubChem 2023 update. Nucleic Acids Research 51, D1373–D1380. URL: https://academic.oup.com/nar/article/51/D1/D1373/6777787, doi:10.1093/nar/gkac956.

Klein, C.S., Marsh, G.D., Petrella, R.J., Rice, C.L., 2003. Muscle fiber number in the biceps brachii muscle of young and old men. Muscle and Nerve 28, 62–68. URL: https://onlinelibrary.wiley.com/doi/10.1002/mus.10386, doi:10.1002/mus.10386.

Koronowski, K.B., Kinouchi, K., Welz, P.S., Smith, J.G., Zinna, V.M., Shi, J., Samad, M., Chen, S., Magnan, C.N., Kinchen, J.M., Li, W., Baldi, P., Benitah, S.A., Sassone-Corsi, P., 2019. Defining the Independence of the Liver Circadian Clock. Cell 177, 1448–1462.e14. URL: https://doi.org/10.1016/j.cell.2019.04.025, doi:10.1016/j.cell.2019.04.025.

Koronowski, K.B., Sassone-Corsi, P., 2021. Communicating clocks shape circadian homeostasis. Science 371. doi:10.1126/science.abd0951.

Kwiecień, I., Soko-lowska, M., W-lodek, L., 2007. Nephroprotective effect of cystathionine is due to its diverse action on the kidney and Ehrlich ascites tumor cells. Pharmacological Reports 59, 553–564. URL: http://www.ncbi.nlm.nih.gov/pubmed/18048956.

Lee, Y.N., Nechushtan, H., Figov, N., Razin, E., 2004. The Function of Lysyl-tRNA Synthetase and Ap4A as Signaling Regulators of MITF Activity in FcϵRI-Activated Mast Cells. Immunity 20, 145–151. URL: https://linkinghub.elsevier.com/retrieve/pii/S1074761304000202, doi:10.1016/S1074-7613(04)00020-2.

Lemons, D.E., Downey, J.A., 1994. Control of the Circulation in the Limbs, in: The Physiological Basis of Rehabilitation Medicine. Elsevier, pp. 365–391. URL: https://linkinghub.elsevier.com/retrieve/pii/B9781483178189500204, doi:10.1016/b978-1-4831-7818-9.50020-4.

Liu, X., Lu, S., Song, K., Shen, Q., Ni, D., Li, Q., He, X., Zhang, H., Wang, Q., Chen, Y., Li, X., Wu, J., Sheng, C., Chen, G., Liu, Y., Lu, X., Zhang, J., 2020. Unraveling allosteric landscapes of allosterome with ASD. Nucleic Acids Research 48. URL: http://mdl.shsmu.edu.cn/ASD, doi:10.1093/nar/gkz958.

Lonsdale, J., Thomas, J., Salvatore, M., Phillips, R., Lo, E., Shad, S., Hasz, R., Walters, G., Garcia, F., Young, N., Foster, B., Moser, M., Karasik, E., Gillard, B., Ramsey, K., Sullivan, S., Bridge, J., Magazine, H., Syron, J., Fleming, J., Siminoff, L., Traino, H., Mosavel, M., Barker, L., Jewell, S., Rohrer, D., Maxim, D., Filkins, D., Harbach, P., Cortadillo, E., Berghuis, B., Turner, L., Hudson, E., Feenstra, K., Sobin, L., Robb, J., Branton, P., Korzeniewski, G., Shive, C., Tabor, D., Qi, L., Groch, K., Nampally, S., Buia, S., Zimmerman, A., Smith, A., Burges, R., Robinson, K., Valentino, K., Bradbury, D., Cosentino, M., Diaz-Mayoral, N., Kennedy, M., Engel, T., Williams, P., Erickson, K., Ardlie, K., Winckler, W., Getz, G., DeLuca, D., Daniel MacArthur, Kellis, M., Thomson, A., Young, T., Gelfand, E., Donovan, M., Meng, Y., Grant, G., Mash, D., Marcus, Y., Basile, M., Liu, J., Zhu, J., Tu, Z., Cox, N.J., Nicolae, D.L., Gamazon, E.R., Im, H.K., Konkashbaev, A., Pritchard, J., Stevens, M., Flutre, T., Wen, X., Dermitzakis, E.T., Lappalainen, T., Guigo, R., Monlong, J., Sammeth, M., Koller, D., Battle, A., Mostafavi, S., McCarthy, M., Rivas, M., Maller, J., Rusyn, I., Nobel, A., Wright, F., Shabalin, A., Feolo, M., Sharopova, N., Sturcke, A., Paschal, J., Anderson, J.M., Wilder, E.L., Derr, L.K., Green, E.D., Struewing, J.P., Temple, G., Volpi, S., Boyer, J.T., Thomson, E.J., Guyer, M.S., Ng, C., Abdallah, A., Colantuoni, D., Insel, T.R., Koester, S.E., A Roger Little, Bender, P.K., Lehner, T., Yao, Y., Compton, C.C., Vaught, J.B., Sawyer, S., Lockhart, N.C., Demchok, J., Moore, H.F., 2013. The Genotype-Tissue Expression (GTEx) project. Technical Report 6. doi:10.1038/ng.2653.

Mahadevan, R., Edwards, J.S., Doyle, F.J., 2002. Dynamic Flux Balance Analysis of diauxic growth in Escherichia coli. Biophysical Journal 83, 1331–1340. doi:10.1016/S0006-3495(02)73903-9.

Mahadevan, R., Schilling, C.H., 2003. The effects of alternate optimal solutions in constraint-based genome-scale metabolic models. Metabolic Engineering 5, 264–276. URL: https://linkinghub.elsevier.com/retrieve/pii/S1096717603000582, doi:10.1016/j.ymben.2003.09.002.

Martins Conde, P., Pfau, T., Pires Pacheco, M., Sauter, T., 2021. A dynamic multi-tissue model to study human metabolism. npj Systems Biology and Applications 7. URL: https://doi.org/10.1038/s41540-020-00159-1, doi:10.1038/s41540-020-00159-1.

Masri, S., Papagiannakopoulos, T., Kinouchi, K., Liu, Y., Cervantes, M., Baldi, P., Jacks, T., Sassone-Corsi, P., 2016. Lung Adenocarcinoma Distally Rewires Hepatic Circadian Homeostasis. Cell 165, 896–909. URL: http://dx.doi.org/10.1016/j.cell.2016.04.039, doi:10.1016/j.cell.2016.04.039.

Murphy, R.M., Larkins, N.T., Mollica, J.P., Beard, N.A., Lamb, G.D., 2009. Calsequestrin content and SERCA determine normal and maximal Ca2+ storage levels in sarcoplasmic reticulum of fast- and slow-twitch fibres of rat. Journal of Physiology 587, 443–460. URL: https://physoc.onlinelibrary.wiley.com/doi/10.1113/jphysiol.2008.163162, doi:10.1113/jphysiol.2008.163162.

Nguyen, M.K., Kao, L., Kurtz, I., 2009. Calculation of the equilibrium pH in a multiple-buffered aqueous solution based on partitioning of proton buffering: A new predictive formula. American Journal of Physiology - Renal Physiology 296, F1521–F1529. URL: https://www.physiology.org/doi/10.1152/ajprenal.90651.2008, doi:10.1152/ajprenal.90651.2008.

Pacheco, M.P., Sauter, T., 2018. The FASTCORE family: For the fast reconstruction of compact context-specific metabolic networks models, in: Methods in Molecular Biology. volume 1716, pp. 101–110. URL: http://link.springer.com/10.1007/978-1-4939-7528-0_4, doi:10.1007/978-1-4939-7528-0_4.

Pagliarini, R., Castello, R., Napolitano, F., Borzone, R., Annunziata, P., Mandrile, G., De Marchi, M., Brunetti-Pierri, N., di Bernardo, D., 2016. In Silico Modeling of Liver Metabolism in a Human Disease Reveals a Key Enzyme for Histidine and Histamine Homeostasis. Cell reports 15, 2292–2300. URL: http://www.ncbi.nlm.nih.gov/pubmed/27239044 http://www.pubmedcentral.nih.gov/articlerender.fcgi?artid=PMC4906368, doi:10.1016/j.celrep.2016.05.014.

Paine, P.L., Moore, L.C., Horowitz, S.B., 1975. Nuclear envelope permeability. Nature 254, 109–114. URL: http://www.ncbi.nlm.nih.gov/pubmed/1117994, doi:10.1038/254109a0.

Palsson, B.O., 2015. Systems biology: Constraint-based reconstruction and analysis. Cambridge University Press. URL: https://www.cambridge.org/core/product/identifier/9781139854610/type/book, doi:10.1017/CBO9781139854610.

Parlakgül, G., Arruda, A.P., Pang, S., Cagampan, E., Min, N., Güney, E., Lee, G.Y., Inouye, K., Hess, H.F., Xu, C.S., Hotamışlıgil, G.S., 2022. Regulation of liver subcellular architecture controls metabolic homeostasis. Nature 603, 736–742. URL: https://www.nature.com/articles/s41586-022-04488-5, doi:10.1038/s41586-022-04488-5.

Pedregosa, F., Varoquaux, G., Gramfort, A., Michel, V., Thirion, B., Grisel, O., Blondel, M., Prettenhofer, P., Weiss, R., Dubourg, V., Vanderplas, J., Passos, A., Cournapeau, D., Brucher, M., Perrot, M., Duchesnay, E., 2011. Scikit-learn: Machine learning in Python. Journal of Machine Learning Research 12, 2825–2830.

Pitzer, K.S., 2018. Activity Coefficients in Electrolyte Solutions. CRC Press. URL: https://www.taylorfrancis.com/books/9781351077927, doi:10.1201/9781351069472.

RDKit,. RDKit: Open-source cheminformatics. URL: https://www.rdkit.org, doi: 10.5281/zenodo.591637.

Reinke, H., Asher, G., 2019. Crosstalk between metabolism and circadian clocks. Nature Reviews Molecular Cell Biology 20, 227–241. URL: https://www.nature.com/articles/s41580-018-0096-9, doi:10.1038/s41580-018-0096-9.

Rimessi, A., Giorgi, C., Pinton, P., Rizzuto, R., 2008. The versatility of mitochondrial calcium signals: From stimulation of cell metabolism to induction of cell death. Biochimica et Biophysica Acta - Bioenergetics 1777, 808–816. URL: https://linkinghub.elsevier.com/retrieve/pii/S0005272808005987, doi:10.1016/j.bbabio.2008.05.449.

Robinson, J.L., Kocabaş, P., Wang, H., Cholley, P.E., Cook, D., Nilsson, A., Anton, M., Ferreira, R., Domenzain, I., Billa, V., Limeta, A., Hedin, A., Gustafsson, J., Kerkhoven, E.J., Svensson, L.T., Palsson, B.O., Mardinoglu, A., Hansson, L., Uhlén, M., Nielsen, J., 2020. An atlas of human metabolism. Science Signaling 13. URL: https://pubmed.ncbi.nlm.nih.gov/32209698/, doi:10.1126/scisignal.aaz1482.

Romani, A.M., 2011. Cellular magnesium homeostasis. Archives of Biochemistry and Biophysics 512, 1–23. URL: https://linkinghub.elsevier.com/retrieve/pii/S0003986111001822, doi:10.1016/j.abb.2011.05.010.

Sanchez, E.J., Lewis, K.M., Danna, B.R., Kang, C.H., 2012. High-capacity Ca 2+ binding of human skeletal calsequestrin. Journal of Biological Chemistry 287, 11592–11601. URL: https://linkinghub.elsevier.com/retrieve/pii/S0021925820480925, doi:10.1074/jbc.M111.335075.

Sehdev., R.D.Z.S.B.J.S., 2023. Physiology, Renal Blood Flow and Filtration. Treasure Island (FL): StatPearls Publishing. URL: https://www.ncbi.nlm.nih.gov/books/NBK482248/.

Seifert, E.L., Ligeti, E., Mayr, J.A., Sondheimer, N., Hajnóczky, G., 2015. The mitochondrial phosphate carrier: Role in oxidative metabolism, calcium handling and mitochondrial disease. Biochemical and Biophysical Research Communications 464, 369–375. URL: https://linkinghub.elsevier.com/retrieve/pii/S0006291X15010335, doi:10.1016/j.bbrc.2015.06.031.

Smith, J.G., Koronowski, K.B., Mortimer, T., Sato, T., Greco, C.M., Petrus, P., Verlande, A., Chen, S., Samad, M., Deyneka, E., Mathur, L., Blazev, R., Molendijk, J., Kumar, A., Deryagin, O., Vaca-Dempere, M., Sica, V., Liu, P., Orlando, V., Parker, B.L., Baldi, P., Welz, P.S., Jang, C., Masri, S., Benitah, S.A., Muñoz-Cánoves, P., Sassone-Corsi, P., 2023. Liver and muscle circadian clocks cooperate to support glucose tolerance in mice. Cell Reports 42, 112588. URL: https://linkinghub.elsevier.com/retrieve/pii/S2211124723005995, doi:10.1016/j.celrep.2023.112588.

Smith, J.G., Koronowski, K.B., Sato, T., Greco, C., Petrus, P., Verlande, A., Chen, S., Samad, M., Deyneka, E., Mathur, L., Blazev, R., Molendijk, J., Mortimer, T., Kumar, A., Deryagin, O., Vaca-Dempere, M., Liu, P., Orlando, V., Parker, B.L., Baldi, P., Welz, P.S., Jang, C., Masri, S., Benitah, S.A., Muñoz-Cánoves, P., Sassone-Corsi, P., 2022. Interrogating Metabolic Interactions Between Skeletal Muscle and Liver Circadian Clocks In Vivo. bioRxiv, 2022.02.27.482160 URL: https://www.biorxiv.org/content/10.1101/2022.02.27.482160v1%0A https://www.biorxiv.org/content/10.1101/2022.02.27.482160v1.abstract, doi:10.1101/2022.02.27.482160.

Snijders, T., Aussieker, T., Holwerda, A., Parise, G., van Loon, L.J., Verdijk, L.B., 2020. The concept of skeletal muscle memory: Evidence from animal and human studies. Acta Physiologica 229. URL: https://onlinelibrary.wiley.com/doi/10.1111/apha.13465, doi:10.1111/apha.13465.

Stewart, P.A., 1978. Independent and dependent variables of acid-base control. Respiration Physiology 33, 9–26. URL: https://linkinghub.elsevier.com/retrieve/pii/0034568778900798, doi:10.1016/0034-5687(78)90079-8.

Stewart, P.A., 1981. How to Understand Acid-Base. A Quantitative Acid-Base Primer for Biology and Medicine.. volume 57. Edward Arnold. doi:10.1086/412691.

Stewart, P.A., 1983. Modern quantitative acid-base chemistry. Canadian Journal of Physiology and Pharmacology 61, 1444–1461. URL: http://www.nrcresearchpress.com/doi/10.1139/y83-207, doi:10.1139/y83-207.

Stroh, A.M., Lynch, C.E., Lester, B.E., Minchev, K., Chambers, T.L., Montenegro, C.F., Martinez, C.C., Fountain, W.A., Trappe, T.A., Trappe, S.W., 2021. Human adipose and skeletal muscle tissue DNA, RNA, and protein content. Journal of Applied Physiology 131, 1370–1379. URL: https://journals.physiology.org/doi/10.1152/japplphysiol.00343.2021, doi:10.1152/japplphysiol.00343.2021.

Subramanian, A., Tamayo, P., Mootha, V.K., Mukherjee, S., Ebert, B.L., Gillette, M.A., Paulovich, A., Pomeroy, S.L., Golub, T.R., Lander, E.S., Mesirov, J.P., 2005. Gene set enrichment analysis: A knowledge-based approach for interpreting genome-wide expression profiles. Proceedings of the National Academy of Sciences of the United States of America 102, 15545–15550. URL: https://pubmed.ncbi.nlm.nih.gov/16199517/www.pnas.orgcgidoi10.1073pnas.0506580102, doi:10.1073/pnas.0506580102.

Thiele, I., Palsson, B., 2010. A protocol for generating a highquality genome-scale metabolic reconstruction. Nature Protocols 5, 93–121. URL: https://www.nature.com/articles/nprot.2009.203, doi:10.1038/nprot.2009.203.

Thiele, I., Sahoo, S., Heinken, A., Hertel, J., Heirendt, L., Aurich, M.K., Fleming, R.M., 2020. Personalized wholebody models integrate metabolism, physiology, and the gut microbiome. Molecular Systems Biology 16, 8982. URL: https://www.embopress.org/doi/10.15252/msb.20198982, doi:10.15252/msb.20198982.

Thiele, I., Swainston, N., Fleming, R.M., Hoppe, A., Sahoo, S., Aurich, M.K., Haraldsdottir, H., Mo, M.L., Rolfsson, O., Stobbe, M.D., Thorleifsson, S.G., Agren, R., Bolling, C., Bordel, S., Chavali, A.K., Dobson, P., Dunn, W.B., Endler, L., Hala, D., Hucka, M., Hull, D., Jameson, D., Jamshidi, N., Jonsson, J.J., Juty, N., Keating, S., Nookaew, I., Le Novere, N., Malys, N., Mazein, A., Papin, J.A., Price, N.D., Selkov, E., Sigurdsson, M.I., Simeonidis, E., Sonnenschein, N., Smallbone, K., Sorokin, A., Van Beek, J.H., Weichart, D., Goryanin, I., Nielsen, J., Westerhoff, H.V., Kell, D.B., Mendes, P., Palsson, B.O., 2013. A community-driven global reconstruction of human metabolism. Nature Biotechnology 31, 419–425. doi:10.1038/nbt.2488.

Wang, Q., Michalak, M., 2020. Calsequestrin. Structure, function, and evolution. Cell Calcium 90, 102242. URL: https://linkinghub.elsevier.com/retrieve/pii/S0143416020300841, doi:10.1016/j.ceca.2020.102242.

Wang, Z.M., Ying, Z., Bosy-Westphal, A., Zhang, J., Schautz, B., Later, W., Heymsfield, S.B., Müller, M.J., 2010. Specific metabolic rates of major organs and tissues across adulthood: Evaluation by mechanistic model of resting energy expenditure. American Journal of Clinical Nutrition 92, 1369–1377. URL: https://linkinghub.elsevier.com/retrieve/pii/S0002916523021421, doi:10.3945/ajcn.2010.29885.

Wasserman, D.H., 2009. Four grams of glucose. American Journal of Physiology - Endocrinology and Metabolism 296, E11–E21. URL: https://www.physiology.org/doi/10.1152/ajpendo.90563.2008, doi:10.1152/ajpendo.90563.2008.

Wishart, D.S., Feunang, Y.D., Marcu, A., Guo, A.C., Liang, K., Vázquez-Fresno, R., Sajed, T., Johnson, D., Li, C., Karu, N., Sayeeda, Z., Lo, E., Assempour, N., Berjanskii, M., Singhal, S., Arndt, D., Liang, Y., Badran, H., Grant, J., Serra-Cayuela, A., Liu, Y., Mandal, R., Neveu, V., Pon, A., Knox, C., Wilson, M., Manach, C., Scalbert, A., 2018. HMDB 4.0: The human metabolome database for 2018. Nucleic Acids Research 46, D608–D617. URL: https://www.nmrml.org, doi:10.1093/nar/gkx1089.

Wu, G., Xie, X., Lu, Z.H., Ledeen, R.W., 2009. Sodium-calcium exchanger complexed with GM1 ganglioside in nuclear membrane transfers calcium from nucleoplasm to endoplasmic reticulum. Proceedings of the National Academy of Sciences of the United States of America 106, 10829–10834. URL: https://pnas.org/doi/full/10.1073/pnas.0903408106, doi:10.1073/pnas.0903408106.

Wu, M., Wu, C., Song, T., Pan, K., Wang, Y., Liu, Z., 2023. Structure and transport mechanism of the human calcium pump SPCA1. Cell Research 33, 533–545. URL: https://www.nature.com/articles/s41422-023-00827-x, doi:10.1038/s41422-023-00827-x.

Xin, H., Zhang, J., Huang, R., Li, L., Lam, S.M., Shui, G., Deng, F., Zhang, Z., Li, M.D., 2022. Circadian signatures of adipose tissue in diet-induced obesity. Frontiers in Physiology 13. URL: https://www.frontiersin.org/articles/10.3389/fphys.2022.953237/full, doi:10.3389/fphys.2022.953237.

Xiong, J., Zhu, M.X., 2016. Regulation of lysosomal ion homeostasis by channels and transporters. Science China Life Sciences 59, 777–791. URL: http://link.springer.com/10.1007/s11427-016-5090-x, doi:10.1007/s11427-016-5090-x.

Xu, Z., Zhang, D., He, X., Huang, Y., Shao, H., 2016. Transport of Calcium Ions into Mitochondria. Current Genomics 17, 215–219. URL: http://www.eurekaselect.com/openurl/content.php?genre=article&issn=1389-2029&volume=17&issue=3&spage=215, doi:10.2174/1389202917666160202215748.

Young, S., Egginton, S., 2009. Allometry of skeletal muscle fine structure allows maintenance of aerobic capacity during ontogenetic growth. Journal of Experimental Biology 212, 3564–3575. URL: https://journals.biologists.com/jeb/article/212/21/3564/19009/Allometry-of-skeletal-muscle-fine-structure-allows, doi:10.1242/jeb.029512.

Zhao, S., Fung-Leung, W.P., Bittner, A., Ngo, K., Liu, X., 2014. Comparison of RNA-Seq and microarray in transcriptome profiling of activated T cells. PLoS ONE 9, e78644. URL: https://dx.plos.org/10.1371/journal.pone.0078644, doi:10.1371/journal.pone.0078644.

